# Lower intake of animal-based products links to improved weight status, independent of depressive symptoms and personality in the general population

**DOI:** 10.1101/2020.02.09.940460

**Authors:** Evelyn Medawar, Cornelia Enzenbach, Susanne Röhr, Arno Villringer, Steffi G. Riedel-Heller, A. Veronica Witte

**Author notes:** Corresponding author: Evelyn Medawar, Max Planck Institute for Human Cognitive and Brain Sciences, Stephanstraße 1a, 04103 Leipzig, Germany. Preregistered analysis plan on OSF https://osf.io/4w69q.

## Abstract

**Background:** Restricting animal-based products from diet may exert beneficial effects on weight status, however whether this is also true for emotional health is unclear. Moreover, differential personality traits may underlie restrictive eating habits and therefore potentially confound diet-health associations. To systematically assess whether restrictive dietary intake of animal-based products relates to lower weight and higher depressive symptoms, and how this is linked to personality traits in the general population.

**Methods:** Cross-sectional data was taken from the baseline LIFE-Adult study collected from 2011-2014 in Leipzig, Germany (n = 8943). Main outcomes of interest were 12-month dietary frequency of animal-derived products measured using a Food Frequency Questionnaire (FFQ), body mass index (BMI) (kg/m^2^), and the Center of Epidemiological Studies Depression Scale (CES-D). Personality traits were assessed in a subsample of n = 7906 using the Five Factor Inventory (NEO-FFI).

**Findings:** Higher restriction of animal-based product intake was associated with a lower BMI (age-, sex- and education-adjusted, n = 8943; ß = −.07, p < .001), but not depression score. Personality, i.e. lower extraversion (F _(1,7897)_ = 9.8, p = .002), was related to frequency of animal product intake. Further, not diet but personality was significantly associated with depression, i.e. higher neuroticism (ß = .024), lower extraversion (ß = −.006), lower agreeableness (ß = −.001), lower conscientiousness (ß = −.007) and higher BMI (ß = .004) (all p < .001, overall model, R^2^ = .21). The beneficial association with lower weight seemed to be driven by the frequency of meat product intake and not secondary animal products. Likewise, the overall number of excluded food items from the individual diet was associated with a lower BMI (age-, sex- and education-adjusted, n = 8938, ß = −.15, p < .001) and additionally with lower depression scores (ß = −.004, t = −4.1, p < .001, R^2^ = .05, corrected for age, sex and education), also when additionally correcting for differences in personality traits (ß = −.003, t = −2.7, p = .007, R^2^ = .21).

**Interpretation:** Higher restriction of animal-based products in the diet was significantly associated with a lower BMI, but not with depressive symptoms scores in a large well-characterized population-based sample of adults. In addition, we found that certain personality traits related to restricting animal-based products – and that those traits, but not dietary habits, explained a considerable amount of variance in depressive symptoms. Upcoming longitudinal studies need to confirm these findings and to test the hypothesis if restricting animal-based products, esp. primary animal products ((processed) meat, wurst), conveys benefits on weights status, hinting to a beneficial relationship of animal-based restricted diets in regard to prevention and treatment of overweight and obesity.

**Funding:** We thank all study participants. We very much appreciate the help of the physicians who performed the clinical examinations and data collection, in particular Ulrike Scharrer, Annett Wiedemann, Kerstin Wirkner and her team. This work was supported by LIFE—Leipzig Research Centre for Civilisation Diseases, University of Leipzig. LIFE is funded by means of the European Union, by means of the European Social Fund (ESF), by the European Regional Development Fund (ERDF), and by means of the Free State of Saxony within the framework of the excellence initiative. This work was supported by a scholarship (EM) by the German Federal Environmental Foundation and by the grants of the German Research Foundation contract grant number CRC 1052 “Obesity mechanisms” Project A1 (AV) and WI 3342/3-1 (AVW). The corresponding author had full access to all the data in the included in the analysis and had final responsibility for the decision to submit for publication.

**Graphical abstract:** 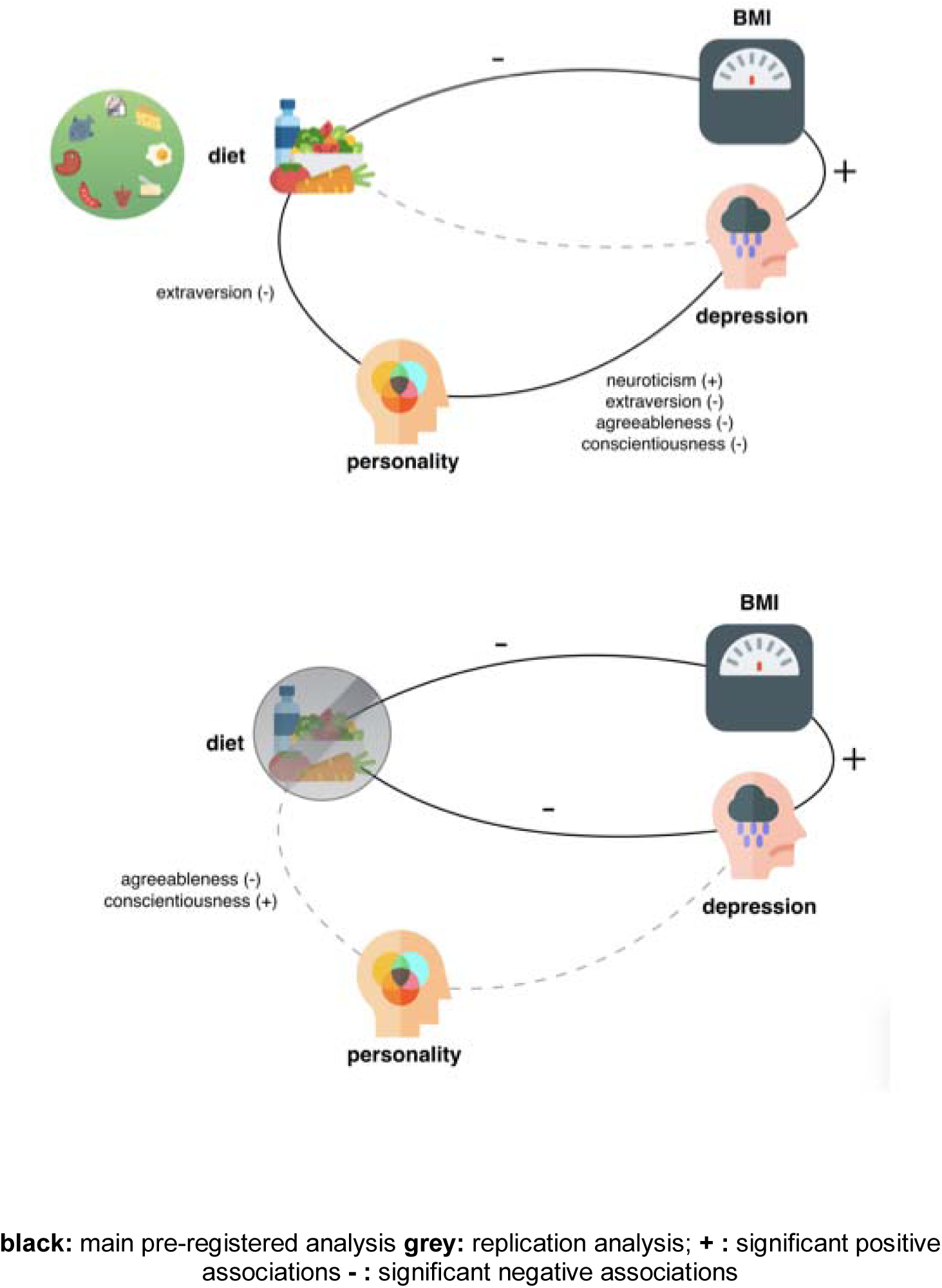

## Research in context

### Evidence before this study

Restriction of animal-based products in eating patterns such as vegetarian and vegan diets are widely debated to convey either health benefits or risks. Large population-based studies as well as randomized controlled trials investigated medium to long-term effects of plant-based diets on different aspects of health, i.e. metabolic and mental health, with partly inconsistent findings. Yet, recent evidence accumulates indicating that benefits of plant-based diets are multi-fold and affect both human health and planetary health in a positive way.

### Added value of this study

To our knowledge, no previous study has combined all three domains of diet, metabolic health and mental health. Here, we aim to assess these domains in a comprehensive manner in order to understand the complex interplay of lifestyle factors (such as diet) on health-related measures. Moreover, we extended the analysis to further lifestyle-relevant measures, such as personality traits, which we could show to be a (strong) confounding factor for the association observed between the restriction of animal-based products and depression. Our analyses for the first time include a large population-based sample of German individuals investigating dietary patterns on a continuous scale on their association with weight status, personality traits and depressive symptoms.

### Implications of all the available evidence

Our analysis contributes to the public health relevance of restricting animal-based products by showing beneficial effects on weight status without impeding personality traits or depressive symptoms. Our results emphasize the relevance of reducing frequency of animal-based products for health reasons for the general population, supporting the adoption of a flexitarian, meat-reduced diet.

## Introduction

Animal product-restrictive eating patterns such as vegetarian and vegan diets are debated to convey either health benefits or risks (reviewed in ^1^). For example, epidemiological studies like the Adventist studies (n = 22,000-96,000) found that plant-based eating habits compared to omnivorous diets are associated with lower all-cause mortality and less frequent with cardiovascular diseases ^2,3^. Other studies like the EPIC-Oxford study (n~64,000) ^4^ and the „45 and Up Study” (n~267,000) ^5^ showed however no effect of a plant-based diet on mortality rate. The 18 years follow-up of the EPIC-Oxford study showed a decrease of ischaemic heart disease prevalence on the one hand, and an increased odds ratio for total stroke on the other hand in fish-eaters and vegetarians compared to meat-eaters ^6^. Intervention studies in small to moderate sample sizes (n∼ 100) indicated that medium-term (12-74 weeks) vegan diets, compared to omnivorous diets, leads to weight loss and to a decrease in type 2 diabetes symptoms, even when caloric intake was comparably low between the diets ^7–9^.

While the exact mechanisms mediating these effects are far from fully understood, improved energy metabolism, reductions of systemic low-grade inflammation as well as changes in microbiome-gut-brain signaling might play a pivotal role ^1,10–14^.

Further, individuals showing restrictive eating patterns, i.e. excluding animal-derived food, may be more or less prone to develop mood disturbances compared to those with omnivorous eating styles: large epidemiological studies (n = 6,422-90,380) showed higher depressive symptoms in vegetarians and vegans ^15–17^ and in those with orthorexic behaviour ^18^. Yet other (smaller) cross-sectional (n = 620) and interventional (n = 39-291) studies proposed a positive effect of plant-based diets on well-being and subclinical depression scores ^19–22^. Recently, it has been suggested that not meat-restriction per se, but the number of excluded food groups predicts higher depressive scores ^17^.

In addition, both weight gain and weight loss may relate to depressive symptoms ^23^, and obesity and depression are assumed to share not only certain symptoms but also genetic pathways and personality traits, in particular neuroticism (reviewed in ^24^). For example, studies showed that higher neuroticism and lower conscientiousness correlate with a higher BMI and more depressive symptoms ^25,26^. Moreover, differences in personality traits and demographic factors such as age, sex and education have also been linked to more or less restrictive lifestyle habits, including diet ^27–29^.

Taken together, these factors likely introduce confounding in studies assessing the relationship between diet, weight status and depressive symptoms separately. However, these complex dependencies have not always been taken into account in previous studies, rendering a definitive conclusion on whether animal product-restrictive eating habits convey health benefits or health risks difficult. We therefore aimed to systematically determine the interplay between animal-restrictive vs. omnivorous dietary habits (measured on a continuum as frequency of animal-based food intake), weight status, depressive symptoms and personality traits in a large population-based sample of adults in Germany.

We hypothesize that: 1) higher restriction of animal-based products is associated with lower BMI (kg/m^2^), even when accounting for potential confounding factors, 2) higher restriction of animal-based products is associated with certain personality traits, measured using the Five-Factor Inventory (NEO-FFI), 3) higher restriction of animal-based products is associated with higher depression scores (measured using CES-D), yet the association may attenuate when taking differences in demographics and personality traits into account.

## Methods

All analyses and hypotheses have been preregistered in the Open Science Framework (OSF) at https://osf.io/4w69q. Participants were drawn from the population-based LIFE-Adult cohort, which aims to explore causes and developments of common civilization diseases such as obesity, depression and dementia (see ^28^ for details). Volunteers underwent anthropometric measurements and answered extensive questionnaires regarding dietary habits, depressive mood and personality (see below for details).

### Inclusion criteria

The initial dataset consisted of n = 10,083 participants taken from the Adult Baseline and Adult Baseline Plus samples. Subjects were included into the analysis if valid and complete measures of age, sex, education, BMI, CES-D and FFQ were available, resulting in a sample of n = 8,943 (sample 1) and a subsample with additional available personality trait measure of n = 7,906 (sample 2, Figure 1). Note that results from sample 2 may slightly deviate from the previously reported pilot analyses in the OSF registration due to partially non-overlapping samples and an extension to a personality questionnaire that was widely available in the dataset.

**Figure 1:**
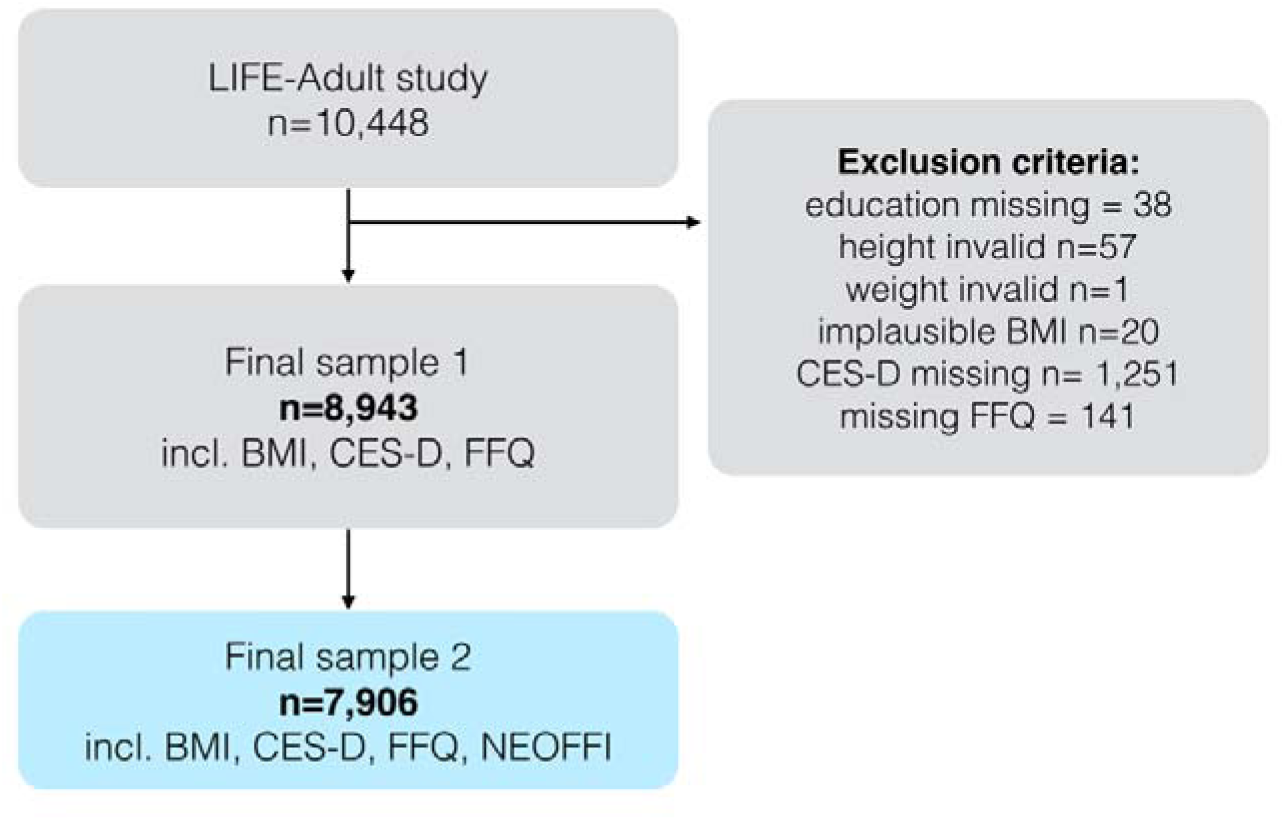
Flowchart of sample selection for sample 1 and sample 2. Abbreviations: BMI=Body-Mass-Index, CES-D=Center of Epidemiological Studies Depression Scale, FFQ=Food Frequency Questionnaire, NEOFFI=NEO Five-Factor-Inventory.

### Ethics

The institutional ethics board of the Medical Faculty of the University of Leipzig raised no concerns regarding the study protocol and all participants provided written informed consent.

### Demographics

Education levels were computed according to Comparative Analysis of Social Mobility in Industrial Nations levels (CASMIN) ^30^ into three levels (low, middle, and high).

### Anthropometry

Body weight was measured with scale SECA 701, height was measured with height rod SECA 220 (SECA Gmbh & Co. KG). Body weight (kg) and body height (m) were used to calculate body-mass-index (BMI) (kg/m^2^). For additional analyses WHO classification for obesity was used: underweight <18.5kg/m^2^, normal-weight >=18.5 and <25kg/m^2^, overweight >=25 and <30kg/m^2^, obese >=30kg/m^2^.

### Personality

Personality traits were measured with the German version of the Big Five via Short Forms (16-Adjective Measure) ^31^; subscales computed for Neuroticism, Extraversion, Openness, Agreeableness and Conscientiousness by building summed scores according to the test’s manual. In a subsample personality traits were measured with the German version of the NEOFFI-30 ^32,33^.

### Depressive scores

Depressive scores (self-reported) were assessed by the Centre of Epidemiologic Studies-Depression (CES-D) scale ^34^.

### Dietary restriction scores (DRS)

Food group items were taken from a questionnaire asking for self-reported food intake frequency over the last 12 months. A composite score for the restriction of animal-derived food items was calculated (Figure 2), including 9 questions regarding the following food groups: meat, processed meat, wurst, fish, eggs, dairy (yoghurt and cream cheese), cheese, milk and butter (animal DRS). Answers ranged from *multiple times daily* (1 per item; 9 for summed score), *daily/(almost) daily, multiple times a week, weekly, 2-3 times monthly, 1 or less a month* to *(almost) never* (7 per item; 63 for summed score). The higher the score, the lower the frequency of consumption of animal-based products. Light products were recoded from 1-5 to 1-7, and either the normal or the light product was chosen for scoring depending on higher frequency; if both were equally frequent, the normal item was chosen (applicable for wurst, dairy, cheese, butter and milk). Measures were ordinal, but for analysis purposes treated as linear, which is a common procedure for scoring lifestyle questionnaire data ^35,36^ and has been shown to perform robustly in parametric analyses (discussed in ^37^). Note that the questionnaire did not include an option such as “I prefer not to answer” or “I don’t know”. Missing values were replaced by the population mean in line with recommendation to use imputation for missing values in nutritional epidemiology ^38^. Subjects with >20% of missing answers out of the 33 food items (excl. drink items) were excluded from the analysis (code and supplementary info available here (https://osf.io/m7hxk/?view_only=91863f44bae44371a1317072334df9fd).

**Figure 2:**
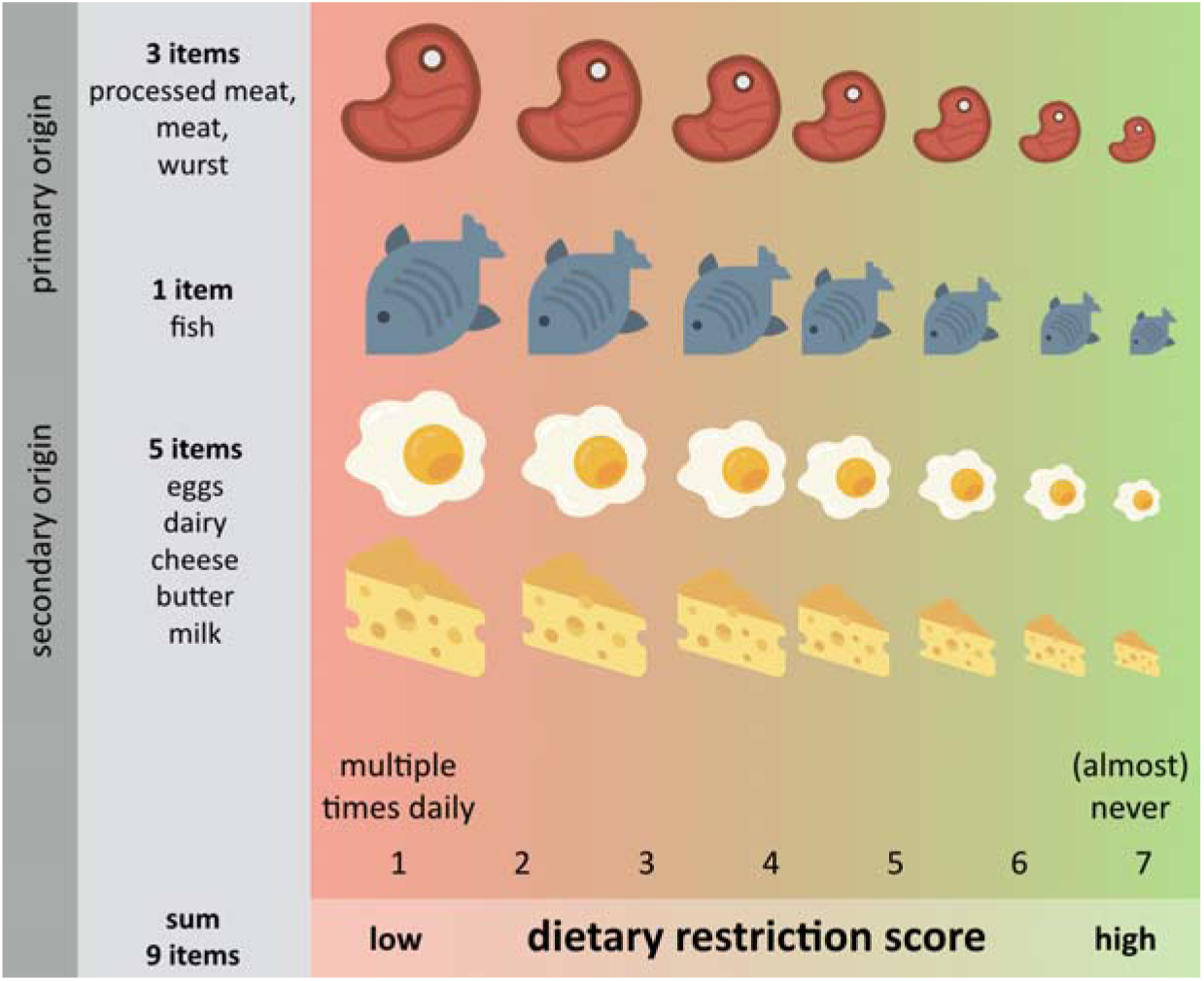
Concept of dietary restriction score (DRS) based on the frequency of consumption of animal-based products over the last 12 months based on 9 items from the FFQ. Copyright icons: all icons by Smashicons.

To further investigate the difference between leaving out primary (meat, bone, and marrow, representing meat-restrictive diets) and/or secondary (stemming from animal labor like milk, representing vegetarian diets) animal products from the diet, we further tested whether potential associations were specific to either food groups by computing two additional scores a) primary DRS and b) secondary DRS (Suppl. Table 1).

An additional score represents the number of restricted food items (adapted from ^17^ by counting all *(almost) never* items of 33 items FFQ (excluding drinks and light products) (score min. 0 to max. 33) within the last 12 months (5.1±2.9 items (mean±SD), range 0-19) (overall DRS).

All computed scores were normally distributed (skewness < 1.0, kurtosis <= 2.0) (Suppl. Figure 1). Moderate positive correlations were observed between meat and wurst (ρ = .46), processed meat and meat (ρ = .26), processed meat and wurst (ρ = .22), dairy and cheese (ρ = .42), and dairy and milk (ρ = .28) consumption (Suppl. Figure 2).

### Statistical models

The main analysis included linear regression models to examine the association of animal DRS and BMI (model 1), depressive symptoms (model 3) and personality traits in a multivariate analysis of covariance (MANCOVA) (model 2). More specifically, model 1 tested whether animal DRS predicted BMI, adjusting for age, sex and education. Model 2 tested whether animal DRS (factor) was associated with the different personality traits (five subscales of the NEO-FFI as dependent variables), accounting for age, sex and education (covariates). Model 3 tested whether animal DRS predicted CES-D when accounting for age, sex and education; and additionally accounting for personality factors and BMI. All variables were normally distributed (skewness < |1.06|, kurtosis < |2.08|), personality traits (skewness < |1.05|, kurtosis < |3.2|), except for CES-D (skewness 1.4, kurtosis = 3.3), which was therefore log-transformed (log10(CES-D+1). Analyses were computed in R version 3.6.1 using lm, lm.beta and ggplot2 for visualization. Statistical significance was set at alpha = 0.05/3 = 0.015 in the main analyses to adjust for multiple testing with the Bonferroni method and at p < 0.05 in all additional analyses.

## Results

We included 8,943 subjects for analyses regarding diet, BMI and depressive symptoms (see Table 1 for demographics), and 7,906 participants in sample 2 for the subsample analysis additionally investigating personality traits (see Table 2).

**Table 1:**
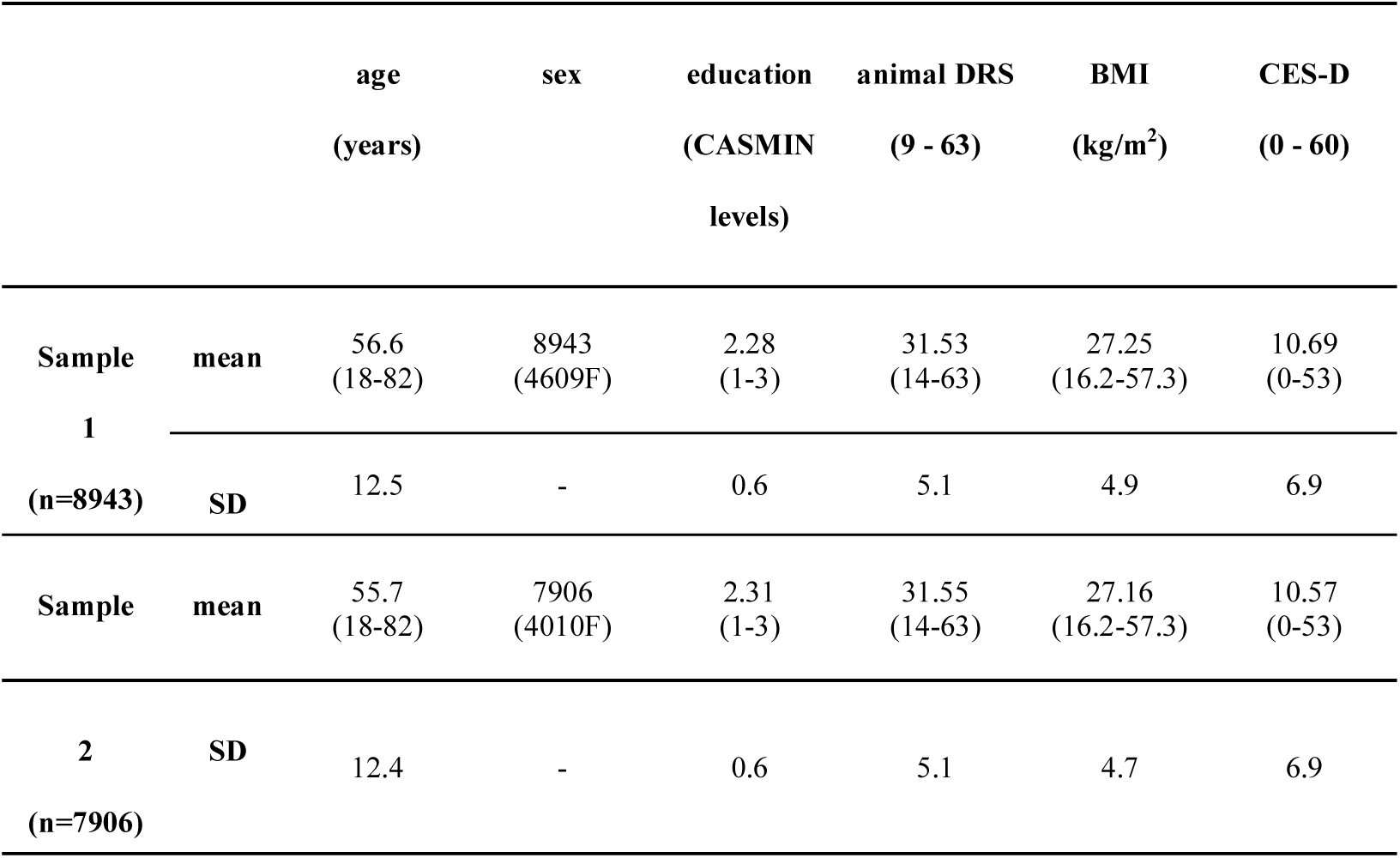
Demographic characteristics for sample 1 and sample 2.

**Table 2:**
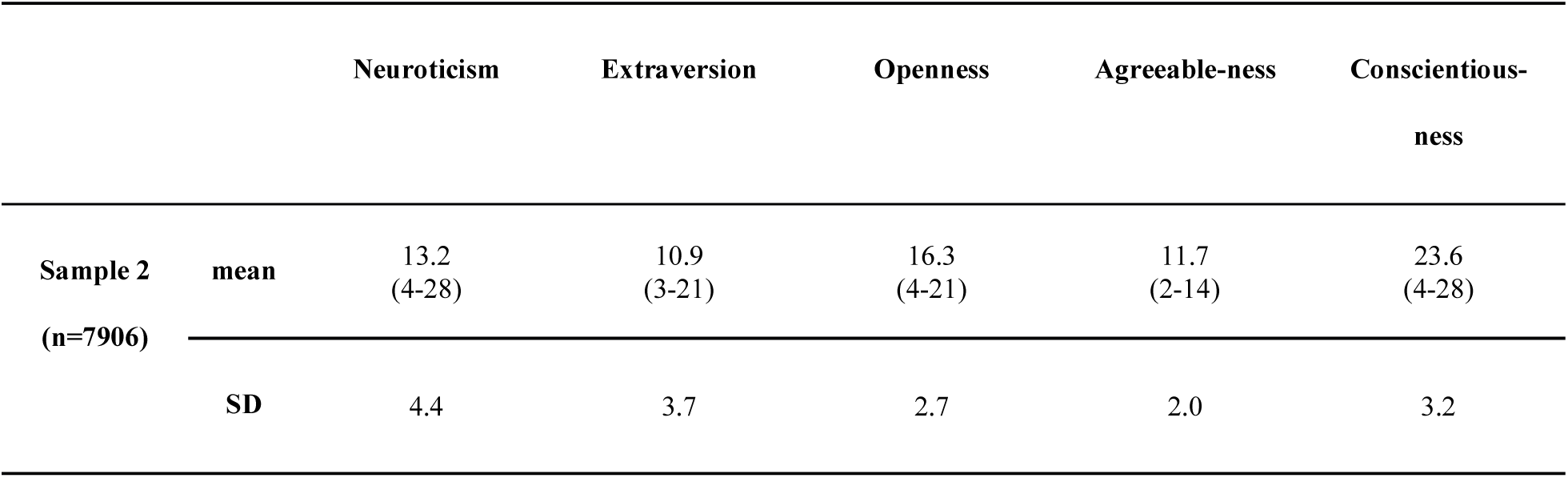
Personality traits according to the five factor personality questionnaire NEO-FFI (16 items) for sample 2 (n=7,906).

Linear regression models detected that lower animal DRS, i.e. higher frequency of animal-based products consumption, related to higher BMI in sample 1 (n = 8943; ß = −.07, p < .001), corrected for confounders (age, sex, education). Higher age, being male and lower education were also significantly associated with higher BMI, with the four factors together explaining about 6% of the variance in BMI (overall model adj. R^2^ = .06, p < .001) (Figure 3A, Table 3). Here, age showed the steepest slope (n = 8943; ß = .08, p < .001; Figure 3B). Similar results emerged when restricting the analysis to the smaller sample 2 (data not shown). When additionally correcting for personality traits the association between BMI and animal DRS remains significant (n = 7906; ß = −.07, p < .001), further certain personality traits show significant associations with BMI (neuroticism: ß = −.05, p < .001; openness: ß = −.05, p < .02; agreeableness: ß = .13, p < .001; conscientiousness: ß = −.2, p < .001; all n = 7906) (Table 3).

**Figure 3:**
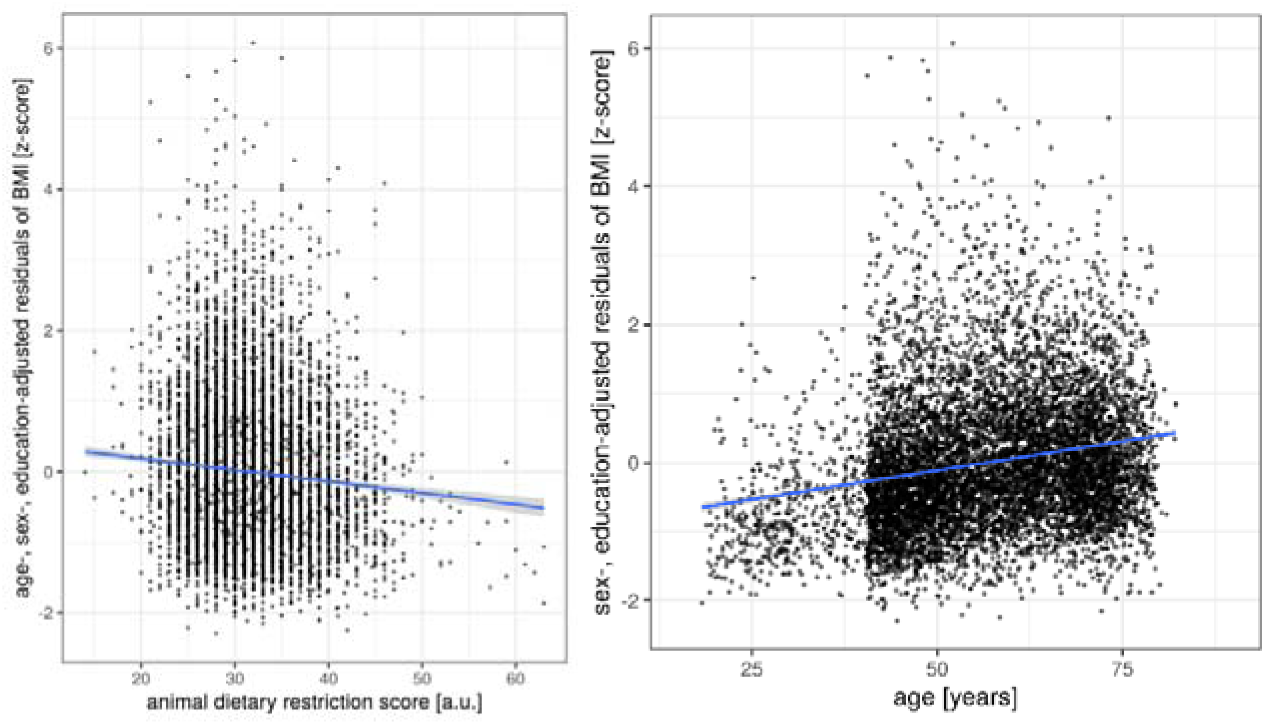
Association between BMI and demographic and lifestyle factors A) animal DRS B) age, residuals plotted according to regression model 1 (sample 1 n = 8943). Line gives regression fit. Point size = 1.

**Table 3:** Multiple regression analyses predicting BMI as function of age, sex, education and frequency of animal-based products (n = 8943).

|  | Adj. R2 | B | C.I. | beta | p |
| --- | --- | --- | --- | --- | --- |
| **BMI (model 1)** |  |  |  |  |  |
| Model | 0.06 |  |  |  | <0.001 |
| sex |  | -0.59 | [-0.79 -0.40] | -0.06 | <0.001 |
| education |  | -0.67 | [-0.83 -0.50] | -0.08 | <0.001 |
| age |  | 0.08 | [0.07 0.09] | 0.21 | <0.001 |
| animal DRS |  | -0.07 | [-0.09 -0.05] | -0.06 | <0.001 |
| **BMI (model 1) – sample 2 (df = 7896), corrected for personality** |  |  |  |  |  |
| Model | 0.08 |  |  |  | <0.001 |
| sex |  | -0.55 | [-0.78 -0.33] | -0.06 | <0.001 |
| education |  | -0.65 | [-0.83 -0.47] | -0.08 | <0.001 |
| age |  | 0.09 | [0.09 0.10] | 0.24 | <0.001 |
| animal DRS |  | -0.07 | [-0.09 -0.05] | -0.07 | <0.001 |
| **neuroticism** |  | -0.05 | [-0.08 -0.03] | -0.05 | **0.001** |
| extraversion |  | 0.01 | [-0.02 0.04] | 0.01 | 0.42 |
| **openness** |  | -0.05 | [-0.10 -0.01] | -0.03 | **0.01** |
| **agreeableness** |  | 0.13 | [0.07 0.19] | 0.05 | **<0.001** |
| **conscientiousness** |  | -0.20 | [-0.23 -0.16] | -0.13 | **<0.001** |

*B/beta represent unstandardized/standardized regression coefficients*

Further, in sample 2 we found a significant association between frequency of animal-based products and personality traits, when correcting for age, sex and education (n = 7906; MANCOVA, F _(5,7897)_ = 2.8, p < .02): higher restriction of animal products was negatively associated with extraversion (F _(1,7897)_ = 9.8, p = .002) (Figure 4, Table 4). Although non-significant, animal DRS was positively associated with neuroticism (F _(1,7897)_ = 3.5, p = .06) and negatively with openness (F _(1,7897)_ = 3.4, p = .07). Likewise, sex was significantly associated with all five personality traits; age and education with four of them (all except for agreeableness) (Table 4).

**Figure 4:**
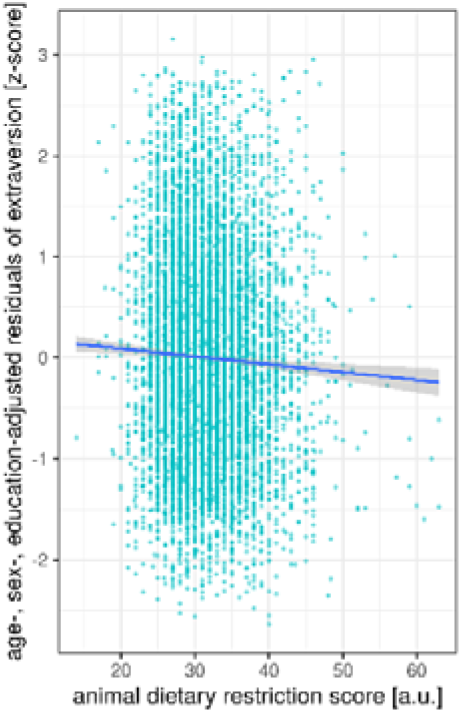
Association between animal DRS and extraversion, residuals plotted according to regression model 2 (sample 1 n = 8943). Line gives regression fit. Point size = 1.

**Table 4:** MANCOVA analysis of animal DRS, age, sex, education on personality (n = 7906).

|  | Pillai's trace | F | df | num df | den df | p |
| --- | --- | --- | --- | --- | --- | --- |
| **NEOFFI (model 2) (all factors, corrected for age, sex, education)** |  |  |  |  |  |  |
| **sex** | 0.17 | 322.2 | 1 | 5 | 7897 | **<0.001** |
| **education** | 0.04 | 66.9 | 1 | 5 | 7897 | **<0.001** |
| **age** | 0.04 | 69.3 | 1 | 5 | 7897 | **<0.001** |
| **animal DRS** | 0.002 | 2.8 | 1 | 5 | 7897 | **0.016** |
| **NEOFFI Neuroticism** |  |  |  |  |  |  |
| **sex** |  | 327.6 | 1 | 5 | 7897 | **<0.001** |
| **education** |  | 113.5 | 1 | 5 | 7897 | **<0.001** |
| **age** |  | 28.5 | 1 | 5 | 7897 | **<0.001** |
| animal DRS |  | 3.5 | 1 | 5 | 7897 | 0.06 |
| **NEOFFI Extraversion** |  |  |  |  |  |  |
| **sex** |  | 15.9 | 1 | 5 | 7897 | **<0.001** |
| **education** |  | 71.1 | 1 | 5 | 7897 | **<0.001** |
| **age** |  | 152.7 | 1 | 5 | 7897 | **<0.001** |
| **animal DRS** |  | 9.8 | 1 | 5 | 7897 | **0.002** |
| **NEOFFI Openness** |  |  |  |  |  |  |
| **sex** |  | 7.3 | 1 | 5 | 7897 | **0.007** |
| **education** |  | 208.4 | 1 | 5 | 7897 | **<0.001** |
| **age** |  | 4.6 | 1 | 5 | 7897 | **0.03** |
| animal DRS |  | 3.4 | 1 | 5 | 7897 | 0.07 |
| **NEOFFI Agreeableness** |  |  |  |  |  |  |
| **sex** |  | 953.5 | 1 | 5 | 7897 | **<0.001** |
| education |  | 1.0 | 1 | 5 | 7897 | 0.33 |
| age |  | 0.7 | 1 | 5 | 7897 | 0.39 |
| animal DRS |  | 0.03 | 1 | 5 | 7897 | 0.87 |
| **NEOFFI Conscientiousness** |  |  |  |  |  |  |
| **sex** |  | 137.4 | 1 | 5 | 7897 | **<0.001** |
| **education** |  | 10.7 | 1 | 5 | 7897 | **0.001** |
| **age** |  | 148.4 | 1 | 5 | 7897 | **<0.001** |
| animal DRS |  | 0.0006 | 1 | 5 | 7897 | 0.98 |

Lastly, frequency of animal-based products did not predict variance in depressive symptoms in sample 1 (n = 8943, ß = .001, p = .12), according to a linear regression model (model 3) that corrected for age, sex, and education (overall model: R^2^ = .04, p < .001) (Table 5). This was also the case for sample 2 (n = 7906, animal DRS: ß = .001, p = .10; overall model; R = .04; p < .001), also when additionally correcting for personality traits and BMI (n = 7906, animal DRS: ß = .013, p = .2; overall model; R = .21; p < .001) (Table 5). Instead, higher neuroticism (ß = .4, p < .001), lower extraversion (ß = −.08, p < .001), lower openness (ß = −.07, p < .001), lower conscientiousness (ß = −.08, p < .001) and higher BMI (ß = .06, p < .001) correlated with depressive symptoms (overall model explaining 21% of variance on depression symptom score) (Figure 5, Table 5).

**Figure 5:**
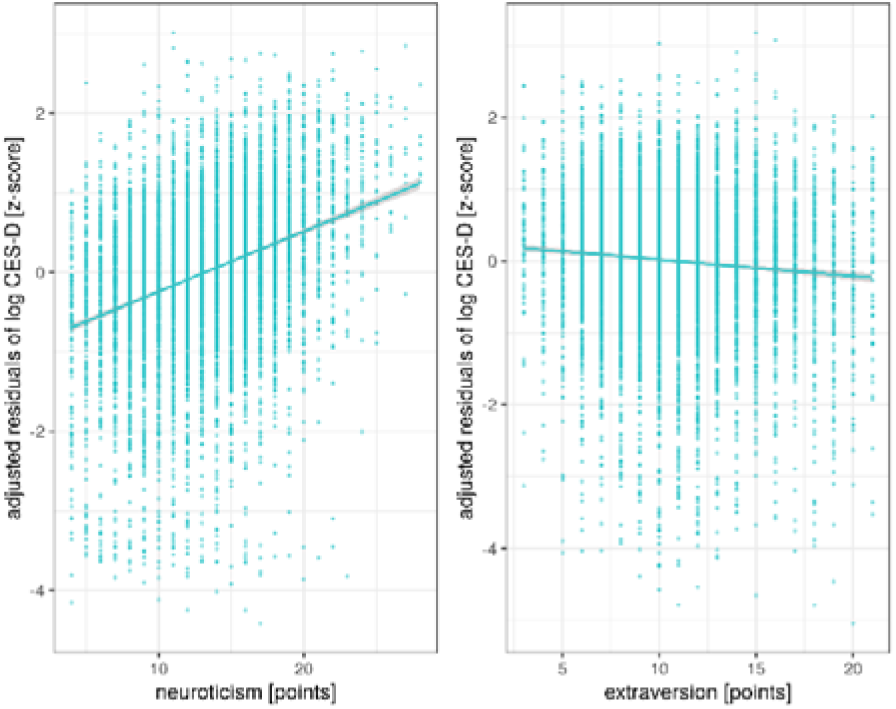
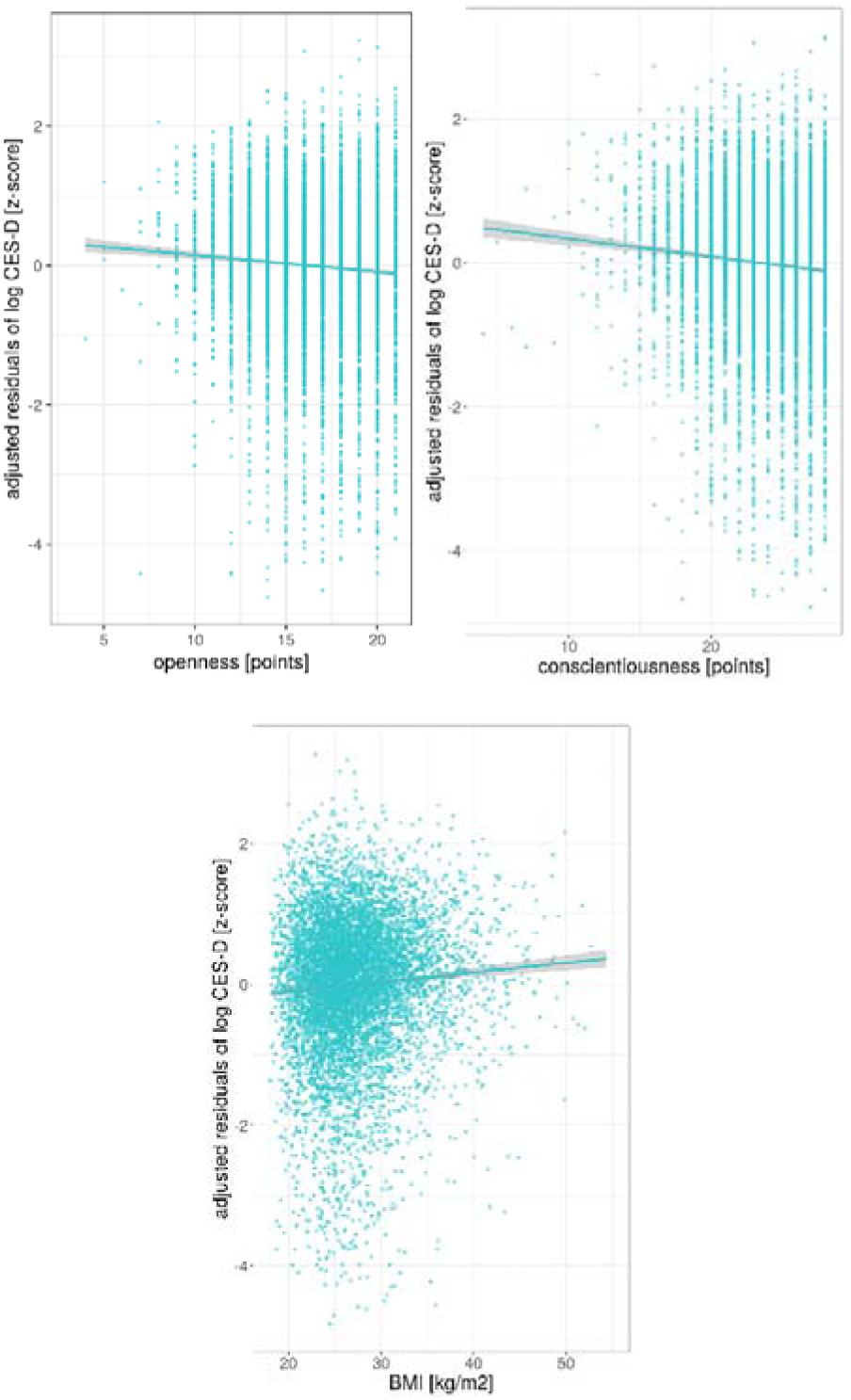
Significant association between personality traits and depressive symptoms in sample 2 (n = 7906) corrected for age, sex, education, animal DRS and the respective four other subscales of personality for neuroticism, extraversion, agreeableness, conscientiousness and BMI. Lines give regression fit. Position size = 2 (for personality) and 1 (BMI).

**Table 5:** Multiple regression analyses predicting CES-D as a function of age, sex, education animal DRS (sample 1, n=8493) and additionally personality traits (sample 2, n = 7906) and BMI.

|  | Adj. R2 | B | C.I. | beta | p |
| --- | --- | --- | --- | --- | --- |
| **CES-D (model 3) - sample 1 (df = 8938)** |  |  |  |  |  |
| Model | 0.04 |  |  |  | <0.001 |
| **sex** |  | 0.04 | [0.029 0.051] | 0.071 | **<0.001** |
| **education** |  | -0.09 | [-0.10 -0.08] | -0.184 | **<0.001** |
| **age** |  | 0.001 | [0.0007 0.0016] | 0.050 | **<0.001** |
| animal DRS |  | 0.001 | [-0.0002 0.0020] | 0.016 | 0.12 |
| **CES-D (model 3) - sample 2 (df = 7901)** |  |  |  |  |  |
| Model | 0.04 |  |  |  |  |
| **sex** |  | 0.04 | [0.0273 0.0523] | 0.069 | **<0.001** |
| **education** |  | -0.09 | [-0.1001 -0.0786] | -0.180 | **<0.001** |
| **age** |  | 0.001 | [0.0006 0.0016] | 0.049 | **<0.001** |
| animal DRS |  | 0.001 | [-0.0002 0.0022] | 0.018 | 0.10 |
| **CES-D (model 3) - sample 2 (df = 7896), corrected for personality** |  |  |  |  |  |
| Model | 0.21 |  |  |  |  |
| sex |  | 0.011 | [-0.001 0.024] | 0.02 | 0.08 |
| **education** |  | -0.06 | [-0.07 -0.05] | -0.12 | **<0.001** |
| **age** |  | 0.0006 | [0.0001 0.0011] | 0.03 | **0.015** |
| animal DRS |  | 0.0005 | [-0.0006 0.0015] | 0.009 | 0.40 |
| **neuroticism** |  | 0.024 | [0.022 0.025] | 0.36 | **<0.001** |
| **extraversion** |  | -0.006 | [-0.008 -0.005] | -0.08 | **<0.001** |
| **openness** |  | -0.007 | [-0.010 -0.005] | -0.07 | **<0.001** |
| agreeableness |  | -0.0004 | [-0.004 0.003] | -0.003 | 0.80 |
| **conscientiousness** |  | -0.008 | [-0.009 -0.006] | -0.08 | **<0.001** |
| **CES-D (model 3) - sample 2 (df = 7895), corrected for personality and BMI** |  |  |  |  |  |
| Model | 0.21 |  |  |  | <0.001 |
| **sex** |  | 0.013 | [0.0008 0.026] | 0.02 | **0.04** |
| **education** |  | -0.06 | [-0.082 -0.039] | -0.11 | **<0.001** |
| age |  | 0.0002 | [-0.066 -0.046] | 0.01 | 0.32 |
| animal DRS |  | 0.001 | [-0.004 0.002] | 0.013 | 0.20 |
| **neuroticism** |  | 0.024 | [0.022 0.025] | 0.36 | **<0.001** |
| **extraversion** |  | -0.006 | [-0.008 -0.005] | -0.08 | **<0.001** |
| **openness** |  | -0.007 | [-0.010 -0.005] | -0.07 | **0.14** |
| agreeableness |  | -0.0009 | [-0.004 0.003] | -0.006 | 0.60 |
| **conscientiousness** |  | -0.007 | [-0.009 -0.005] | -0.08 | **<0.001** |
| **BMI** |  | 0.004 | [ 0.002 0.005] | 0.06 | **<0.001** |

*B/beta represent unstandardized/standardized regression coefficients*

To confirm whether results were not driven by extreme cases with pathological underweight, we excluded underweight individuals (BMI <= 18.5kg/m^2^) from the analysis (n = 51, 17.8±0.6 kg/m^2^ (mean±SD), range 16-18.5). This did not change the results from the main analyses (data not shown).

### Exploratory analyses

Restricting primary animal source products (i.e. (processed) meat, wurst) was significantly associated with a lower BMI (n = 8943; ß = −.25, p < .001, Figure 6), but not restricting intake of secondary animal products (cheese, milk, eggs) (n = 8943, ß = −.02, p = .16) (Table 6). Note the somewhat stronger association of primary animal-based products with BMI compared to the “comprehensive” animal-product DRS score, resulting in a more negative ß coefficient.

**Figure 6:**
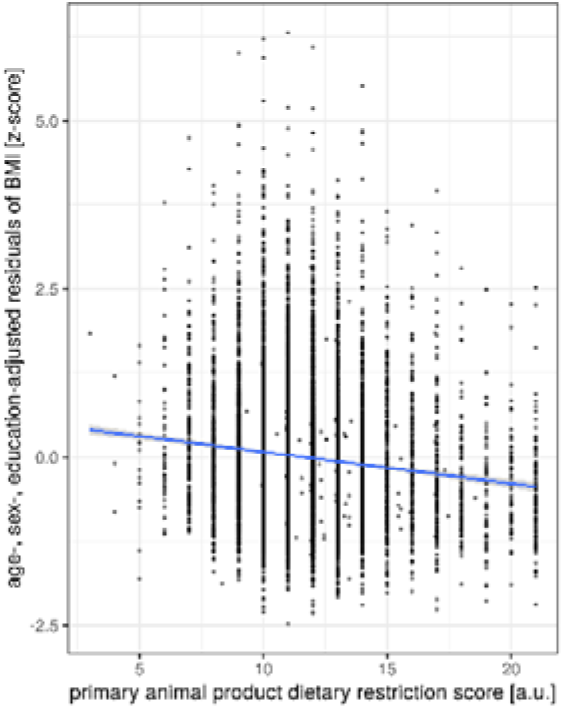
Restrictive animal-based product intake associated with lower BMI. Lines give regression fit. Position size = 1.

**Table 6:**
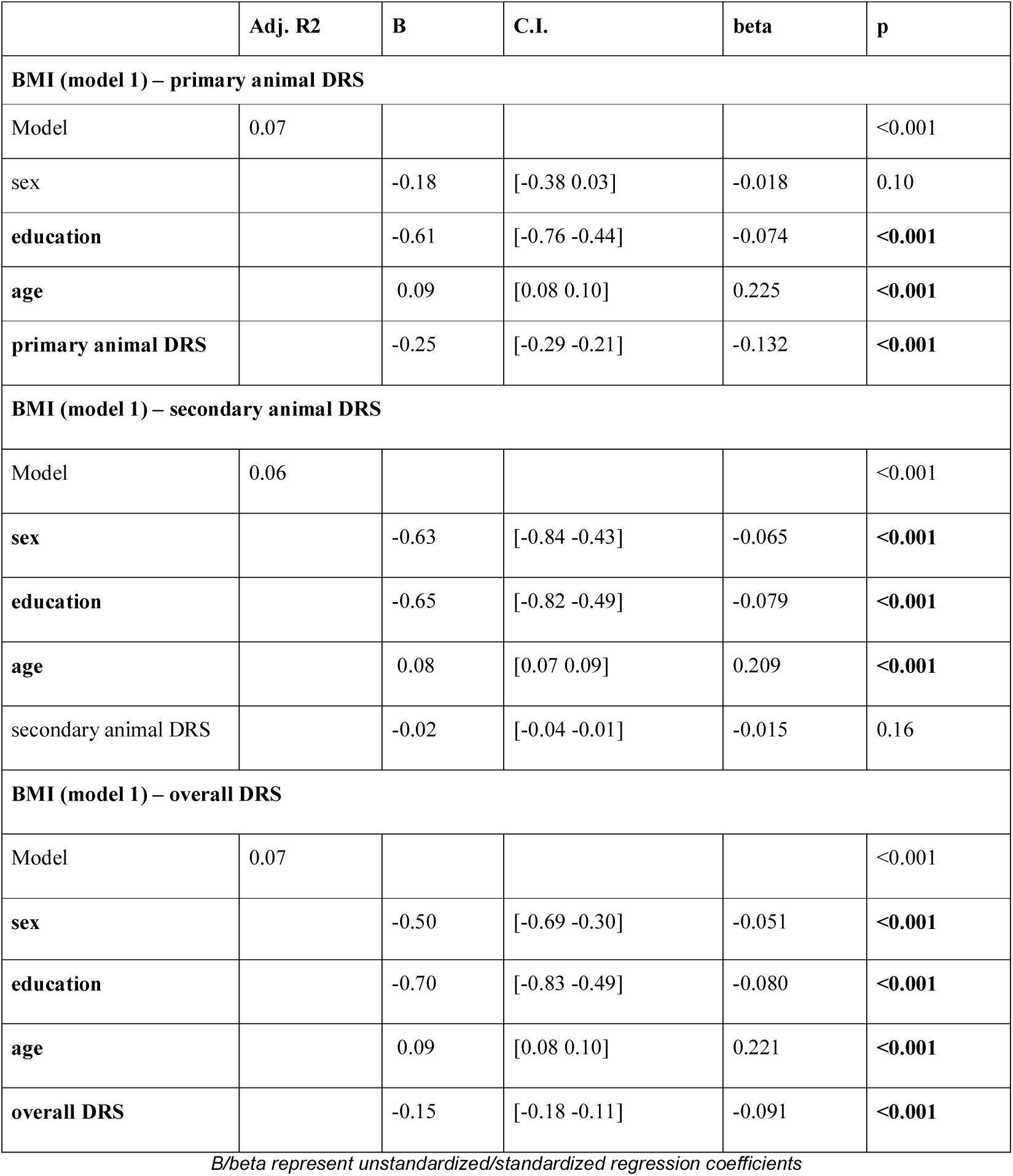
Multiple regression analyses predicting BMI as a function of age, sex, education and restriction of different dietary items (sample 1, n=8493).

|  | Adj. R2 | B | C.I. | beta | p |
| --- | --- | --- | --- | --- | --- |
| **BMI (model 1) – primary animal DRS** |  |  |  |  |  |
| Model | 0.07 |  |  |  | <0.001 |
| sex |  | -0.18 | [-0.38 0.03] | -0.018 | 0.10 |
| **education** |  | -0.61 | [-0.76 -0.44] | -0.074 | **<0.001** |
| **age** |  | 0.09 | [0.08 0.10] | 0.225 | **<0.001** |
| **primary animal DRS** |  | -0.25 | [-0.29 -0.21] | -0.132 | **<0.001** |
| **BMI (model 1) – secondary animal DRS** |  |  |  |  |  |
| Model | 0.06 |  |  |  | <0.001 |
| **sex** |  | -0.63 | [-0.84 -0.43] | -0.065 | **<0.001** |
| **education** |  | -0.65 | [-0.82 -0.49] | -0.079 | **<0.001** |
| **age** |  | 0.08 | [0.07 0.09] | 0.209 | **<0.001** |
| secondary animal DRS |  | -0.02 | [-0.04 -0.01] | -0.015 | 0.16 |
| **BMI (model 1) – overall DRS** |  |  |  |  |  |
| Model | 0.07 |  |  |  | <0.001 |
| **sex** |  | -0.50 | [-0.69 -0.30] | -0.051 | **<0.001** |
| **education** |  | -0.70 | [-0.83 -0.49] | -0.080 | **<0.001** |
| **age** |  | 0.09 | [0.08 0.10] | 0.221 | **<0.001** |
| **overall DRS** |  | -0.15 | [-0.18 -0.11] | -0.091 | **<0.001** |

*B/beta represent unstandardized/standardized regression coefficients*

Investigating differences in personality, higher primary animal DRS was significantly associated with lower neuroticism (F _(1,7897)_ = 27.5, p < .001), higher openness (F _(1,7897)_ = 45.1, p < .001), higher agreeableness (F _(1,7897)_ = 262.5, p < .001) and higher conscientiousness (F _(1,7897)_ = 63.1, p < .001). Higher secondary animal DRS was significantly associated with lower extraversion (F _(1,7897)_ = 11.1, p < .001), lower openness (F _(1,7897)_ = 26.9, p < .001), lower agreeableness (F _(1,7897)_ = 106.7, p < .001) and lower conscientiousness (F _(1,7897)_ = 14.2, p < .001) (all: n = 7906, MANCOVA, corrected for age, sex and education) (Suppl. Figure 4).

In contrast to the comprehensive animal product DRS, the scores displaying restriction of either primary or secondary origin animal products were also associated with lower and higher depression scores, respectively (n = 8943, primary animal-product DRS: ß = −.003, p = .04; secondary animal-product DRS: ß = .002, p = .02; models adjusted for age, sex and education). These divergent associations however failed to reach significance when additionally correcting for personality traits (n = 7906, all |ß| < .002, all p > .10, adjusted for age, sex, education and personality) (Table 7).

**Table 7:** Multiple regression analyses predicting CES-D as a function of age, sex, education and primary and secondary dietary restriction score (sample 1 n = 8943, sample 2 n = 7906).

|  | **Adj. R2** | **B** | **C.I.** | **beta** | **p** |
| --- | --- | --- | --- | --- | --- |
| **CES-D - sample 1 (df = 8938)** |  |  |  |  |  |
| **Model** | 0.04 |  |  |  | **<0.001** |
| **sex** |  | 0.05 | [0.031 0.058] | 0.08 | **<0.001** |
| **education** |  | -0.09 | [-0.100 -0.078] | -0.18 | **<0.001** |
| **age** |  | 0.001 | [0.0007 0.0017] | 0.05 | **<0.001** |
| **primary DRS** |  | -0.003 | [-0.005 -0.00008] | -0.02 | **0.04** |
| **CES-D - sample 2 (df = 7896), corrected for personality** |  |  |  |  |  |
| **Model** | 0.21 |  |  |  | **<0.001** |
| **sex** |  | 0.014 | [0.0008 0.0270] | 0.02 | **0.04** |
| **education** |  | -0.06 | [-0.068 -0.048] | -0.12 | **<0.001** |
| **age** |  | 0.0006 | [0.0001 0.0011] | 0.03 | **0.01** |
| primary DRS |  | -0.002 | [-0.004 -0.001] | -0.01 | 0.21 |
| **neuroticism** |  | 0.024 | [0.022 0.025] | 0.36 | **<0.001** |
| **extraversion** |  | -0.006 | [-0.008 -0.005] | -0.08 | **<0.001** |
| **openness** |  | -0.007 | [-0.010 -0.005] | -0.07 | **<0.001** |
| agreeableness |  | -0.0003 | [-0.004 0.003] | -0.002 | 0.84 |
| **conscientiousness** |  | -0.007 | [-0.009 -0.006] | -0.08 | **<0.001** |
| **CES-D - sample 1 (df = 8938)** |  |  |  |  |  |
| **Model** | 0.04 |  |  |  | **<0.001** |
| **sex** |  | 0.04 | [0.032 0.055] | 0.08 | **<0.001** |
| **education** |  | -0.09 | [-0.10 -0.08] | -0.20 | **<0.001** |
| **age** |  | 0.001 | [0.0007 0.0016] | 0.05 | **<0.001** |
| **secondary DRS** |  | 0.002 | [0.0003 0.003] | -0.03 | **0.02** |
| **CES-D - sample 2 (df = 7896), corrected for personality** |  |  |  |  |  |
| **Model** | 0.21 |  |  |  | **<0.001** |
| **sex** |  | 0.013 | [0.0010 0.0261] | 0.02 | **0.05** |
| **education** |  | -0.06 | [-0.068 -0.048] | -0.12 | **<0.001** |
| **age** |  | 0.0006 | [0.0001 0.0011] | 0.03 | **0.01** |
| secondary DRS |  | 0.001 | [-0.005 0.002] | 0.01 | 0.20 |
| **neuroticism** |  | 0.024 | [0.022 0.025] | 0.36 | **<0.001** |
| **extraversion** |  | -0.006 | [-0.008 -0.005] | -0.08 | **<0.001** |
| **openness** |  | -0.007 | [-0.010 -0.005] | -0.07 | **<0.001** |
| agreeableness |  | -0.0003 | [-0.004 0.003] | -0.002 | 0.84 |
| **conscientiousness** |  | -0.008 | [-0.009 -0.006] | -0.08 | **<0.001** |

Further, we found a strong positive correlation between the frequency of animal-based products (animal DRS) and the number of restricted food groups considering all 33 items (overall DRS) (ρ(8941) = .52, p < .001) (Figure 7A).

**Figure 7:**
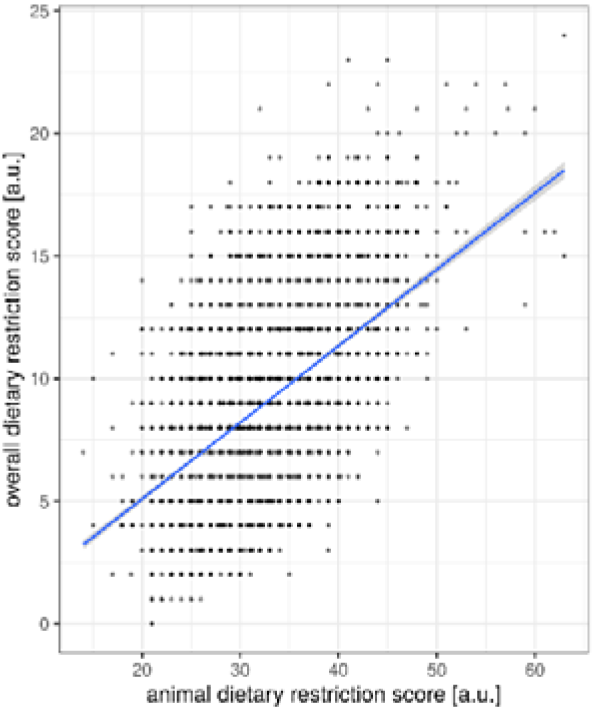
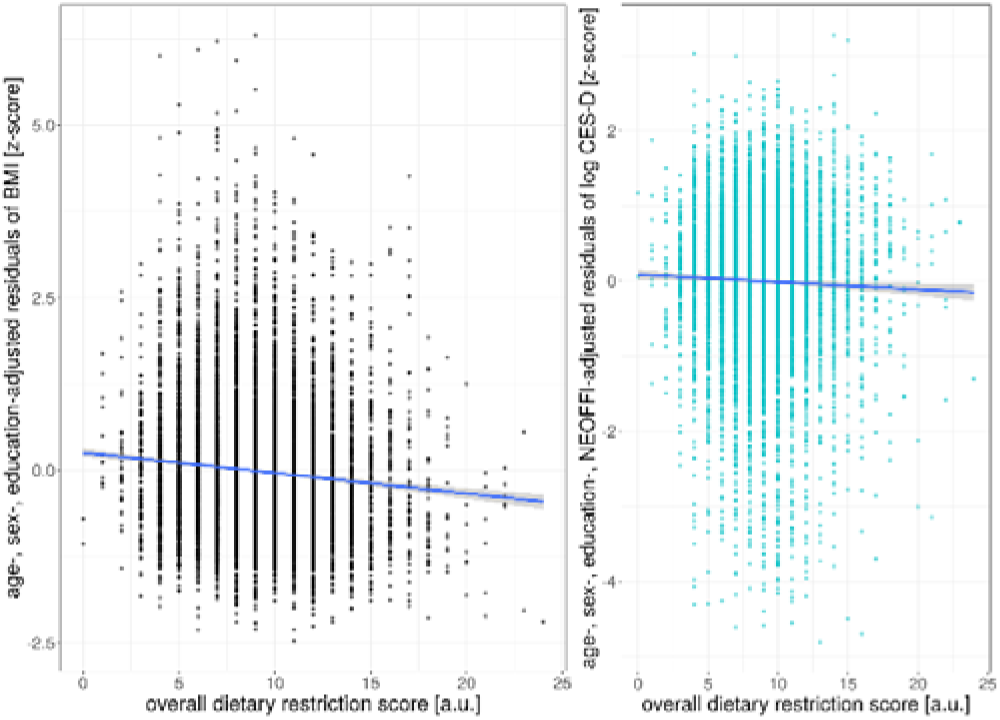
A) Positive association between decreasing frequency of animal-based products and number of excluded food groups. Negative association between overall dietary restriction score and B) BMI and C) CES-D. Position size = 1.

Considering the number of restrictive food items in general, we found that a higher score of total excluded food items related to lower BMI (sample 1: ß = −.15, t = −8.8, p < .001, R^2^ = .07, corrected for age, sex and education; sample 2 similar results (data not shown)) (Figure 7B, Table 6).

The number of restricted food items was significantly associated with lower agreeableness (F _(1,7897)_ = 15.7, p < .001) and higher conscientiousness (F _(1,7897)_ = 53.9, p < .001) (n = 7906, MANCOVA, F _(5,7897)_ = 11.8, p = < .001, for model comparison against null model, corrected for age, sex and education) (Table 8).

**Table 8:** MANCOVA analysis of dietary restriction, age, sex, education on personality (n = 7906).

|  | Pillai's trace | F | df | num df | den df | p |
| --- | --- | --- | --- | --- | --- | --- |
| **NEOFFI (all factors) – sample 2, corrected for age, sex, education** |  |  |  |  |  |  |
| sex | 0.169 | 320.0 | 1 | 5 | 7897 | <0.001 |
| education | 0.041 | 67.4 | 1 | 5 | 7897 | <0.001 |
| age | 0.040 | 65.2 | 1 | 5 | 7897 | <0.001 |
| overall DRS | 0.007 | 11.8 | 1 | 5 | 7897 | <0.001 |
| **NEOFFI Neuroticism** |  |  |  |  |  |  |
| sex |  | 342.0 | 1 | 5 | 7897 | <0.001 |
| education |  | 114.5 | 1 | 5 | 7897 | <0.001 |
| age |  | 28.9 | 1 | 5 | 7897 | <0.001 |
| overall DRS |  | 0.6 | 1 | 5 | 7897 | 0.44 |
| **NEOFFI Extraversion** |  |  |  |  |  |  |
| sex |  | 14.5 | 1 | 5 | 7897 | <0.001 |
| education |  | 72.6 | 1 | 5 | 7897 | <0.001 |
| age |  | 149.3 | 1 | 5 | 7897 | <0.001 |
| overall DRS |  | 0.3 | 1 | 5 | 7897 | 0.6 |
| **NEOFFI Openness** |  |  |  |  |  |  |
| sex |  | 6.1 | 1 | 5 | 7897 | 0.01 |
| **education** |  | 209.8 | 1 | 5 | 7897 | **<0.001** |
| **age** |  | 4.9 | 1 | 5 | 7897 | **0.03** |
| overall DRS |  | 1.6 | 1 | 5 | 7897 | 0.21 |
| **NEOFFI Agreeableness** |  |  |  |  |  |  |
| **sex** |  | 937.3 | 1 | 5 | 7897 | **<0.001** |
| education |  | 0.9 | 1 | 5 | 7897 | 0.34 |
| age |  | 0.2 | 1 | 5 | 7897 | 0.7 |
| **overall DRS** |  | 15.7 | 1 | 5 | 7897 | **<0.001** |
| **NEOFFI Conscientiousness** |  |  |  |  |  |  |
| **sex** |  | 122.4 | 1 | 5 | 7897 | **<0.001** |
| **education** |  | 10.7 | 1 | 5 | 7897 | **0.001** |
| **age** |  | 130.7 | 1 | 5 | 7897 | **<0.001** |
| **overall DRS** |  | 53.9 | 1 | 5 | 7897 | **<0.001** |

Surprisingly, a higher number of restricted food items was weakly yet significantly associated with lower depression scores (ß = −.004, t = −4.1, p < .001, R^2^ = .05, corrected for age, sex and education) (similar results in sample 2 (data not shown)), also when additionally correcting for differences in personality traits (ß = −.003, t = − 2.7, p < .007, R^2^ = .21) (Figure 7C, Table 9).

**Table 9:** Multiple regression analyses predicting CES-D as a function of age, sex, education and dietary restriction score (sample 1 n = 8943, sample 2 n = 7906).

|  | Adj. R2 | B | C.I. | beta | p |
| --- | --- | --- | --- | --- | --- |
| **CES-D - sample 1 (df = 8938)** |  |  |  |  |  |
| **Model** | 0.05 |  |  |  | **<0.001** |
| **sex** |  | 0.04 | [0.032 0.055] | 0.076 | **<0.001** |
| **education** |  | -0.09 | [-0.100 -0.080] | -0.185 | **<0.001** |
| **age** |  | 0.001 | [0.0008 0.0017] | 0.054 | **<0.001** |
| **overall DRS** |  | -0.004 | [-0.006 -0.002] | -0.043 | **<0.001** |
| **CES-D - sample 2 (df = 7901)** |  |  |  |  |  |
| **Model** | 0.04 |  |  |  | **<0.001** |
| **sex** |  | 0.04 | [0.031 0.056] | 0.075 | **<0.001** |
| **education** |  | -0.09 | [-0.100 -0.080] | -0.180 | **<0.001** |
| **age** |  | 0.001 | [0.0008 0.0017] | 0.054 | **<0.001** |
| **overall DRS** |  | -0.005 | [-0.007 -0.002] | -0.048 | **<0.001** |
| **CES-D - sample 2 (df = 7896), corrected for personality** |  |  |  |  |  |
| **Model** | 0.21 |  |  |  | **<0.001** |
| **sex** |  | 0.014 | [0.0010 0.0261] | 0.02 | **0.04** |
| **education** |  | -0.06 | [-0.068 -0.048] | -0.12 | **<0.001** |
| **age** |  | 0.0007 | [0.0002 0.0011] | 0.03 | **0.007** |
| **overall DRS** |  | -0.003 | [-0.004 -0.001] | -0.03 | **0.007** |
| **neuroticism** |  | 0.024 | [0.022 0.025] | 0.36 | **<0.001** |
| **extraversion** |  | -0.006 | [-0.008 -0.005] | -0.08 | **<0.001** |
| **openness** |  | -0.007 | [-0.010 -0.005] | -0.07 | **<0.001** |
| agreeableness |  | -0.0005 | [-0.004 0.003] | -0.004 | 0.76 |
| **conscientiousness** |  | -0.007 | [-0.009 -0.006] | -0.08 | **<0.001** |

B/beta represent unstandardized/standardized regression coefficients

## Discussion

In this large cross-sectional analysis of ∼ 9000 individuals of the general population, lower frequency of eating animal-based products was significantly associated with lower BMI, even when adjusting for confounding effects of age, sex and education. No significant associations emerged between animal-based products consumption and depressive symptom scores when taking personality into account. Frequency of animal-based product consumption was associated with personality traits, in particular with lower extraversion. Surprisingly, not diet but personality was significantly associated with depression scores.

### Weight status

Our finding that eating meat and dairy products less frequently relates to lower BMI is in line with some, but not all, epidemiological and moderate-term randomized interventional trials which point in this direction too ^1,39,40^. In addition, results remained stable even after adjusting for education, which is a strong predictor of both obesity ^41^ and eating habits ^42^, and when taking inter-individual variance in personality traits into account ^43^. Speculating on possible underlying mechanisms, animal-derived products in general are often denser in calories and in total and saturated fats compared to plant-based foods ^44^. In addition, meat and dairy products are oftentimes consumed as processed food, e.g. wurst, deep-fried meat/fish or high-processed snack products, further augmenting their caloric footprint. Thus, lower caloric intake might underlie the observed link between lower frequency of animal-based product consumption and lower BMI. Moreover, recent observations of changes in the gut microbiome due to diet raise the hypothesis that a different distribution of gut bacteria in plant-based dieters alter the ingestion rate of calories from food ^45^, thereby further limiting caloric intake (or bioavailability). However, while these causal pathways between lower frequency of animal-based product intake leading to lower or sustained body weight seem biologically plausible, the association between lower animal-based product intake and lower weight in our cohort might also be a result of lower body weight leading to less animal-based product intake or unknown shared factors that modulate both weight and diet. Future longitudinal observations and interventional trials are needed to further test the above-described hypothesis or its alternatives.

The positive association between restriction of meat products on weight status and the lack of a significant correlation for secondary animal products found in this study and previously by others ^46–48^ could possibly be explained by a higher proportion of highly processed meat items, leading to higher net energy intake and potentially to higher caloric intake ^49^. Further, ongoing discussions on motivations for following certain diets support the view that restraint eating is not directly linked to vegetarian or vegan diets but more common in flexitarians who restrict meat intake with the goal of weight control, which in contrast is not the most common driver in plant-based dieters ^50^.

While due to the cross-sectional design using self-reported FFQ data, estimates of absolute numbers of the strength of the association between diet and BMI are difficult, our findings may be relevant for public health. Considering that changing a conventional Western omnivorous dietary habit to a more plant-based diet, i.e. avoiding (processed) meat and wurst and limiting dairy, cheese and egg intake, would lead to an increase in animal DRS of 20 points, this would translate into ∼ 1.2 kg/m lower BMI. For someone with a frequent intake of primary and secondary animal-product intake (low animal DRS) this could mean for example reducing all animal-based products from multiple times a day to multiple times a week (“flexitarian diet”) or excluding some animal items altogether (“vegan” or “vegetarian” diet). For a 175 cm tall human this would translate into 4 kg of body weight. If obese (e.g. 100 kg, i.e., BMI = 32.7 kg/m), this would mean a reduction of 4% body weight, if overweight (e.g. 80 kg, BMI = 26.1 kg/m) this would mean a reduction of 5% body weight. As a reduction of 5-10% body weight has been shown to significantly reduce obesity-associated co-morbidities in overweight and obesity ^51–57^, restricting dietary intake of animal-based products may be one way to achieve this weight loss goal, and may help to reduce the societal burden of obesity-related diseases and environmental impact caused by high animal-product diets ^39^. However, these calculations have to be interpreted with caution, as our findings rely on self-reported and cross-sectional data only, and we could not quantify dietary intake with regard to the consumed total amounts of food. Future longitudinal observations and interventional trials are needed.

### Depressive symptoms & personality traits

In contrast to previous large cross-sectional studies ^16,17^ and a prospective study in patients with inflammatory bowel disease ^58^, frequency of animal-derived product consumption did not explain variance in depression symptom scores in the current sample.

Yet, intervention studies showed that a plant-based vegan diet compared to a conventional omnivorous diet reduced anxiety and depression or emotional distress ^19–22^, proposing that restricting animal-based products per se may not affect mental health, but rather exert beneficial effects. Notably, we observed that different personality traits and BMI predicted depressive symptom score, which hints towards shared neurobiological mechanisms with obesity ^23,25^. These shared mechanisms might help to explain previous inconsistent findings of a proposed link between restrictive diets and depression: certain personality traits may increase the probability to restrict certain food groups from diet, such as openness and conscientiousness ^59^. Such a correlative link between personality and restrictive eating, although missing in the current data, would thus also apply for restricting animal-based products and may explain higher depressive symptoms in vegetarians or vegans ^16^. Moreover, sociological studies show that animal-restricted dieters are oftentimes stereotyped with a multitude of biases: detrimental health effects, restrictive lifestyle, sentimentalism, extremism, lower perceived masculinity ^60–62^. Aversion to plant-based dieters could lead to higher social exclusion and depressive symptoms as a result. However, more longitudinal studies tracking newly transformed dieters are needed to clarify if avoiding animal-derived products affects mental health.

Differences in our results compared to previous evidence on personality differences in vegetarians may be due to demographic and societal environmental factors. Personality trait differences in vegetarians were found in a cohort of college students ^15^, which might be different to our sample of the general population, in terms of beliefs, motivation of dietary habits and others. Also geographical or cultural settings may influence differences in the results such as westernized (USA ^15^, Germany (this study)) versus mainly-vegetarian Indian cohorts ^29^, who showed higher conscientiousness. Lastly, the popularity and availability of plant-based dishes is a strong modulator of societal acceptance and demand for those kinds of diets. For instance, increasing the offer from one to two plant-based meals in canteens, led to an increase of 40-80% of plant-based meal purchases, underlining the importance of availability as a strong driver ^63^. Since the interest for plant-based diets has been changing dynamically in the last decade, researches should take period and location into account when comparing studies.

Strengths of our study comprise the large, well-characterized population based cohort enabling us to carefully control for important confounders such as education and personality. Moreover, recent studies and meta-analyses focused specifically on intake of red and processed meat and related health outcomes (see e.g. ^64^), however the distinction of restricting diets to not consume primary (vegetarian) and/or secondary animal-products (vegan) is oftentimes overlooked and therefore a strength of our study.

### Limitations

Firstly, limitations of our study include that the results are based on a cross-sectional study design and therefore cannot explain underlying causalities.

Secondly, our analyses are based on self-reported dietary food record, which do not necessarily reflect actual food intake, however, test-retest reliability is generally of good quality ^65^. Moreover, the FFQ used did not ask for quantity of food intake, which limits the interpretability of the observed effects (for further discussion on possible mechanisms see ^1^). Yet, beside this possible inaccuracy of self-reported food intake, we propose that excluding certain food groups for a timeframe of 12 months presumably is a strong and reliable indicator of actual food intake and exclusion of certain food groups.

### Conclusions

Taken together, using a large cross-sectional analysis we observed that a lower frequency of animal-based products was related to lower BMI, while no link between animal-based products intake and depressive symptoms scores emerged. Thus, our findings may suggest that a lower frequency of animal-based products could be able to convey benefits on weights status, hinting to the capacity of plant-based diets as a potentially relevant target for the intervention of obesity and overweight, in particular by reducing the frequency (and probably the amount) of (especially primary source) animal-based products. Long-term interventional trials are needed to test this hypothesis and to clarify the underlying mechanisms.

## Contribution statement

EM and AVW performed literature search and study conception. EM carried out data analysis and figure design. All authors contributed to data interpretation. All authors read and approved the final manuscript.

## Supplementary Material

**Suppl. Figure 1:**
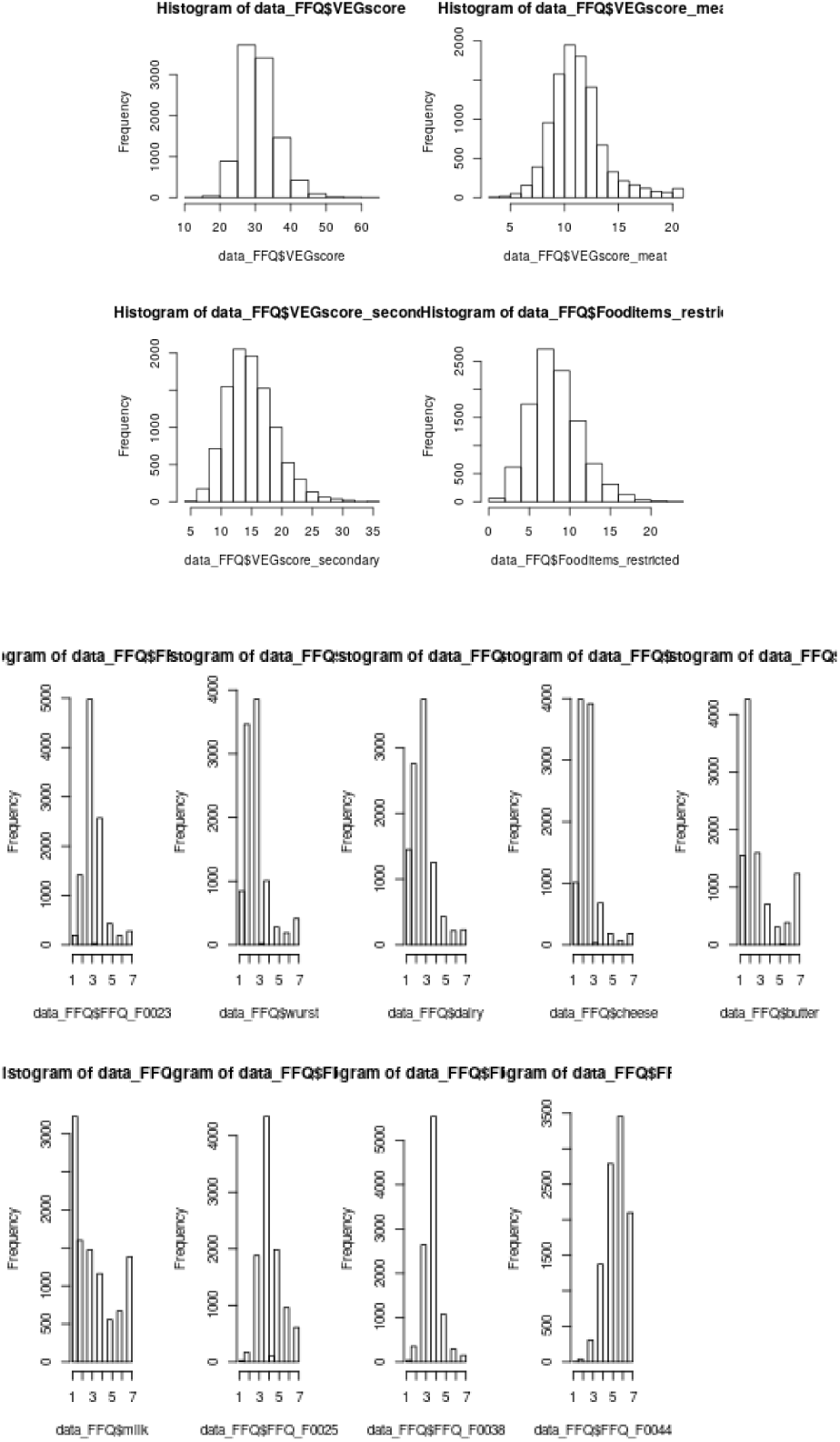
Frequency distribution of the dietary scores. A) animal DRS B) primary animal DRS C) secondary animal DRS and D) overall DRS. All scores are normally distributed (skewness >0.5 and <1). E) Frequency distributions of 9 items used in animal DRS.

**Suppl. Figure 2:**
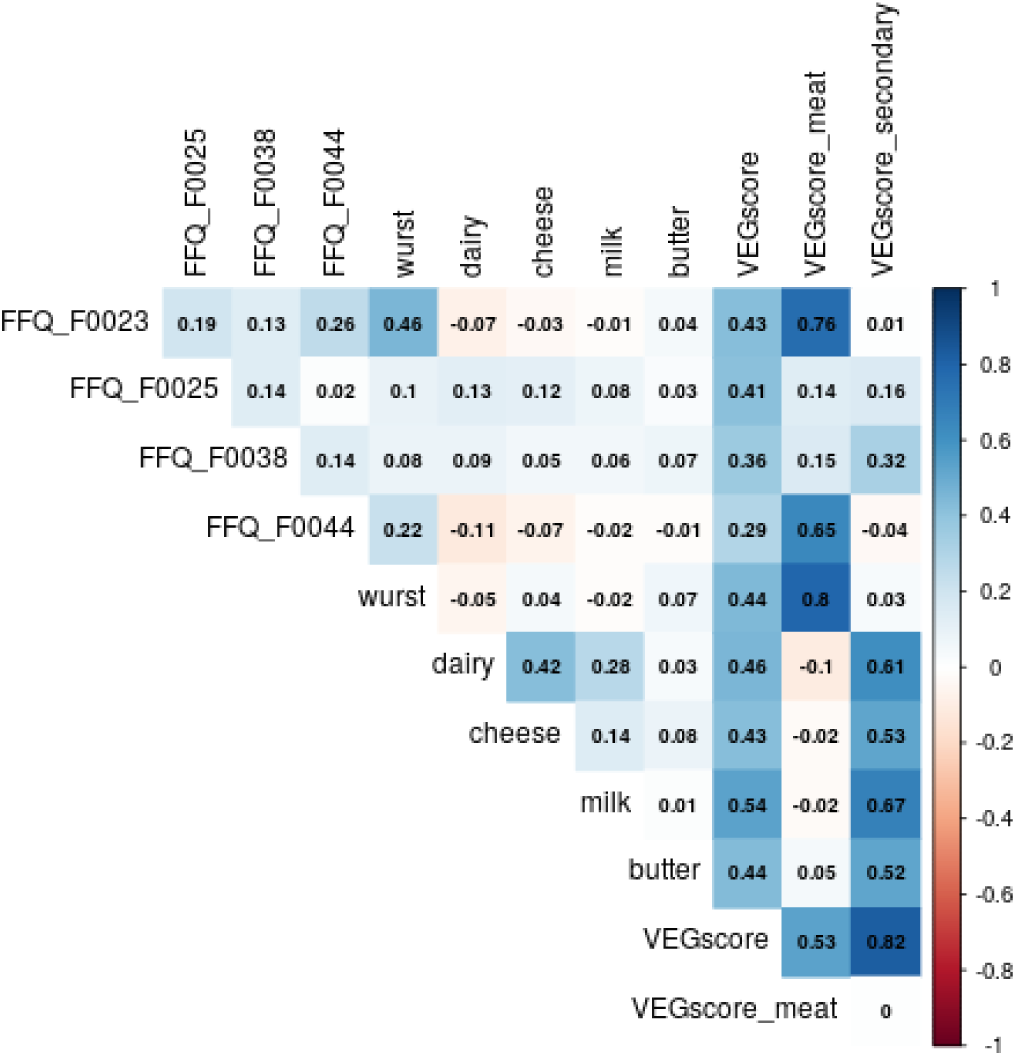
Correlation plot of nine items included in animal DRS.

**Suppl. Figure 3:**
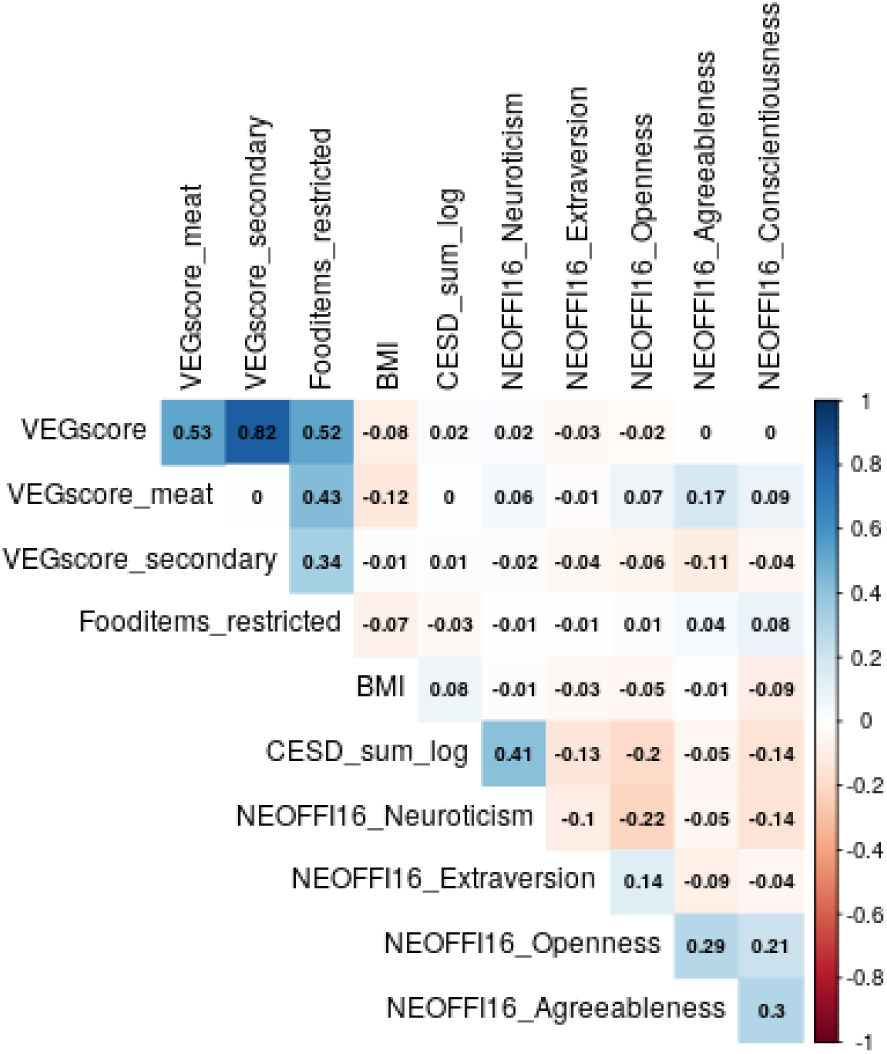
Correlation plot of all measures of interest including dietary patterns, BMI, CES-D and personality traits.

**Suppl. Figure 4:**
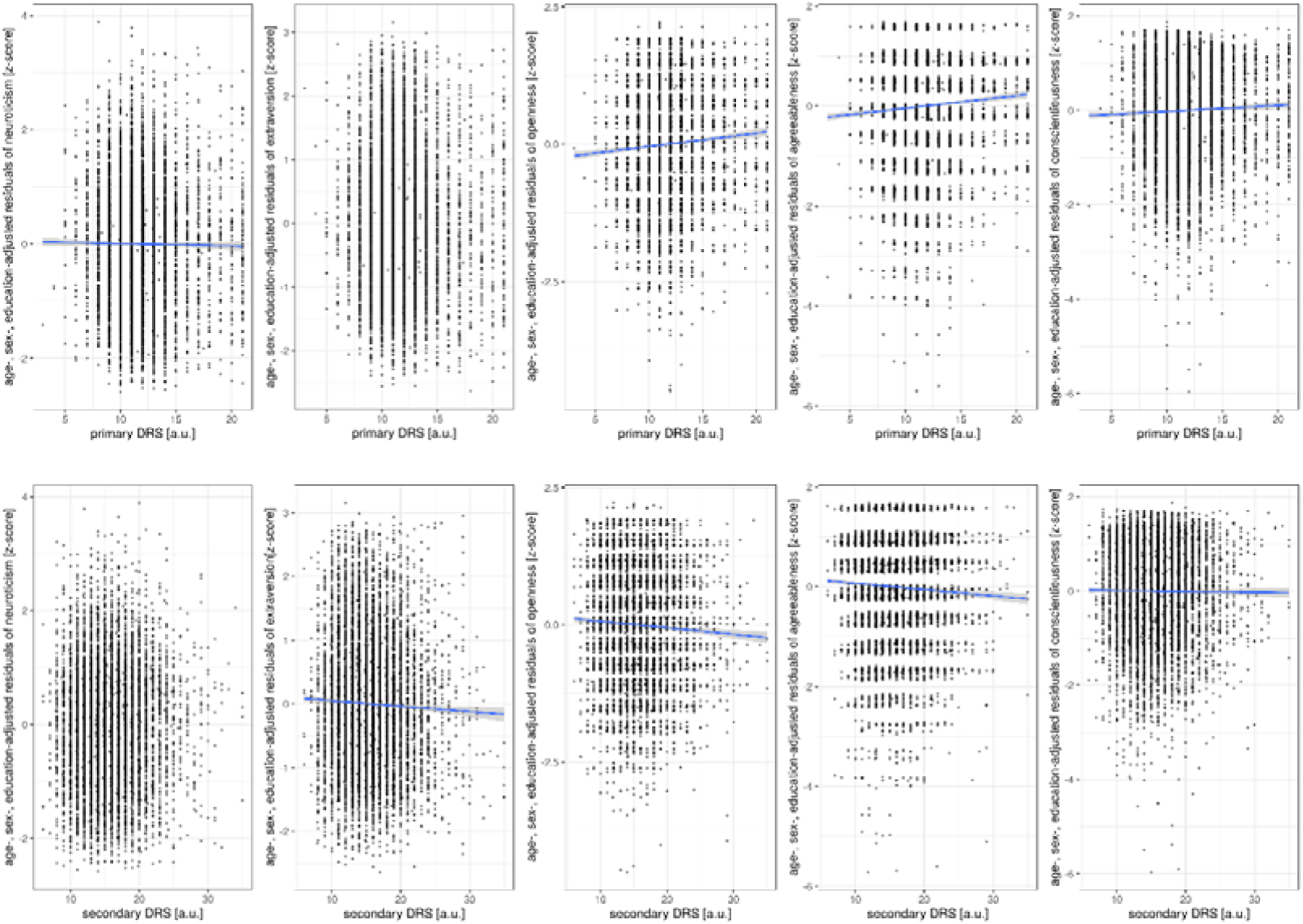
Associations frequency of animal-based products and personality traits (top row: primary DRS; bottom row: secondary DRS).

**Suppl. Table 1:** Summary of computed dietary restriction scores.

|  |  | animal | primary | secondary | overall |
| --- | --- | --- | --- | --- | --- |
|  |  | DRS | animal DRS | animal DRS | DRS |
|  |  | (9 - 63) | (3 - 21) | (5 - 35) | (0 - 33) |
| Sample 1 | mean | 31.5<br>(14-63) | 11.7<br>(3-21) | 15.5<br>(6-35) | 8.7<br>(0-24) |
|  | SD | 5.1 | 2.6 | 4.0 | 3.1 |

## References

1 Medawar E, Huhn S, Villringer A, Veronica Witte A. The effects of plant-based diets on the body and the brain: a systematic review. Transl Psychiatry 2019; 9: 226.

2 Orlich MJ, Singh PN, Sabaté J, et al. Vegetarian dietary patterns and mortality in Adventist Health Study 2. JAMA Intern Med 2013; 173: 1230–8.

3 Le LT, Sabaté J. Beyond meatless, the health effects of vegan diets: Findings from the Adventist cohorts. Nutrients. 2014; 6: 2131–47.

4 Key TJ, Appleby PN, Spencer EA, Travis RC, Roddam AW, Allen NE. Mortality in British vegetarians: results from the European Prospective Investigation into Cancer and Nutrition (EPIC-Oxford)–. Am J Clin Nutr 2009; 89: 1613S–1619S.

5 Mihrshahi S, Ding D, Gale J, Allman-Farinelli M, Banks E, Bauman AE. Vegetarian diet and all-cause mortality: Evidence from a large population-based Australian cohort-the 45 and Up Study. Prev Med (Baltim) 2017; 97: 1–7.

6 Tong TYN, Appleby PN, Bradbury KE, et al. Risks of ischaemic heart disease and stroke in meat eaters, fish eaters, and vegetarians over 18 years of follow-up: results from the prospective EPIC-Oxford study. bmj 2019; 366: l4897.

7 Barnard ND, Cohen J, Jenkins DJA, et al. A low-fat vegan diet and a conventional diabetes diet in the treatment of type 2 diabetes: A randomized, controlled, 74-wk clinical trial. In: American Journal of Clinical Nutrition. 2009. DOI:10.3945/ajcn.2009.26736H.

8 Lee Y-M, Kim S-A, Lee I-K, et al. Effect of a brown rice based vegan diet and conventional diabetic diet on glycemic control of patients with type 2 diabetes: a 12-week randomized clinical trial. PLoS One 2016; 11: e0155918.

9 Kahleova H, Dort S, Holubkov R, Barnard N. A Plant-Based High-Carbohydrate, Low-Fat Diet in Overweight Individuals in a 16-Week Randomized Clinical Trial: The Role of Carbohydrates. Nutrients 2018; 10: 1302.

10 Kahleova H, Fleeman R, Hlozkova A, Holubkov R, Barnard ND. A plant-based diet in overweight individuals in a 16-week randomized clinical trial: metabolic benefits of plant protein. Nutr Diabetes 2018; 8: 58.

11 Kim M, Hwang S, Park E, Bae J. Strict vegetarian diet improves the risk factors associated with metabolic diseases by modulating gut microbiota and reducing intestinal inflammation. Environ Microbiol Rep 2013; 5: 765–75.

12 Tomova A, Bukovsky I, Rembert E, et al. The Effects of Vegetarian and Vegan diets on Gut Microbiota. Front Nutr 2019; 6: 47.

13 Glick-Bauer M, Yeh M-C. The health advantage of a vegan diet: exploring the gut microbiota connection. Nutrients 2014; 6: 4822–38.

14 David LA, Maurice CF, Carmody RN, et al. Diet rapidly and reproducibly alters the human gut microbiome. Nature 2014; 505: 559–63.

15 Forestell CA, Nezlek JB. Vegetarianism, depression, and the five factor model of personality. Ecol Food Nutr 2018; 57: 246–59.

16 Hibbeln JR, Northstone K, Evans J, Golding J. Vegetarian diets and depressive symptoms among men. J Affect Disord 2018; 225: 13–7.

17 Matta J, Czernichow S, Kesse-Guyot E, et al. Depressive Symptoms and Vegetarian Diets: Results from the Constances Cohort. Nutrients 2018; 10: 1695.

18 Luck-Sikorski C, Jung F, Schlosser K, Riedel-Heller SG. Is orthorexic behavior common in the general public? A large representative study in Germany. Eat Weight Disord Anorexia, Bulim Obes 2019; 24: 267–73.

19 Beezhold BL, Johnston CS. Restriction of meat, fish, and poultry in omnivores improves mood: a pilot randomized controlled trial. Nutr J 2012; 11: 9.

20 Kahleova H, Hrachovinova T, Hill M, Pelikanova T. Vegetarian diet in type 2 diabetes– improvement in quality of life, mood and eating behaviour. Diabet Med 2013; 30: 127–9.

21 Agarwal U, Mishra S, Xu J, Levin S, Gonzales J, Barnard ND. A multicenter randomized controlled trial of a nutrition intervention program in a multiethnic adult population in the corporate setting reduces depression and anxiety and improves quality of life: the GEICO study. Am J Heal Promot 2015; 29: 245–54.

22 Beezhold B, Radnitz C, Rinne A, DiMatteo J. Vegans report less stress and anxiety than omnivores. Nutr Neurosci 2015; 18: 289–96.

23 McElroy SL, Kotwal R, Malhotra S, Nelson EB, Keck Jr PE, Nemeroff CB. Are mood disorders and obesity related? A review for the mental health professional. J Clin Psychiatry 2004.

24 Vainik U, Dagher A, Realo A, et al. Personality-obesity associations are driven by narrow traits: A meta-analysis. Obes Rev 2019; 20: 1121–31.

25 Milaneschi Y, Simmons WK, van Rossum EFC, Penninx BWJH. Depression and obesity: evidence of shared biological mechanisms. Mol Psychiatry 2019; 24: 18.

26 Ouakinin SRS, Barreira DP, Gois CJ. Depression and obesity: Integrating the role of stress, neuroendocrine dysfunction and inflammatory pathways. Front Endocrinol (Lausanne) 2018; 9.

27 Allès B, Baudry J, Méjean C, et al. Comparison of sociodemographic and nutritional characteristics between self-reported vegetarians, vegans, and meat-eaters from the Nutrinet-Sante study. Nutrients 2017; 9: 1023.

28 Loeffler M, Engel C, Ahnert P, et al. The LIFE-Adult-Study: objectives and design of a population-based cohort study with 10,000 deeply phenotyped adults in Germany. BMC Public Health 2015; 15: 691.

29 Priyanka, Tanwar K, Kapoor S. Personality Variations of Vegetarian and Non-vegetarian Adolescentss. Indian J Res 2016; 5: 68–70.

30 König W, Lüttinger P, Müller W. A comparative analysis of the development and structure of educational systems: Methodological foundations and the construction of a comparative educational scale. Universität Mannheim, Institut für Sozialwissenschaften, 1988.

31 Herzberg PY, Brähler E. Assessing the Big-Five personality domains via short forms. Eur J Psychol Assess 2006; 22: 139–48.

32 Costa PT, McCrae RR. NEO PI/FFI manual supplement for use with the NEO Personality Inventory and the NEO Five-Factor Inventory. Psychological Assessment Resources, 1989.

33 Körner A, Geyer M, Roth M, et al. Persönlichkeitsdiagnostik mit dem neo-fünf-faktoren-inventar: Die 30-item-kurzversion (neo-ffi-30). PPmP-Psychotherapie· Psychosom Medizinische Psychol 2008; 58: 238–45.

34 Radloff LS. The CES-D scale: A self-report depression scale for research in the general population. Appl Psychol Meas 1977; 1: 385–401.

35 Kurth T, Moore SC, Gaziano JM, et al. Healthy lifestyle and the risk of stroke in women. Arch Intern Med 2006; 166: 1403–9.

36 Flöel A, Witte AV, Lohmann H, et al. Lifestyle and memory in the elderly. Neuroepidemiology 2008; 31: 39–47.

37 Norman G. Likert scales, levels of measurement and the “laws” of statistics. Adv Heal Sci Educ 2010; 15: 625–32.

38 Fraser GE, Yan R, Butler TL, Jaceldo-Siegl K, Beeson WL, Chan J. Missing data in a long food frequency questionnaire: are imputed zeroes correct? Epidemiology 2009; 20: 289.

39 Willett W, Rockström J, Loken B, et al. Food in the Anthropocene: the EAT–Lancet Commission on healthy diets from sustainable food systems. Lancet 2019.

40 Spencer EA, Appleby PN, Davey GK, Key TJ. Diet and body mass index in 38 000 EPIC-Oxford meat-eaters, fish-eaters, vegetarians and vegans. Int J Obes 2003; 27: 728.

41 Wardle J, Waller J, Jarvis MJ. Sex differences in the association of socioeconomic status with obesity. Am J Public Health 2002; 92: 1299–304.

42 Groth M, Fagt S, Brøndsted L. Social determinants of dietary habits in Denmark. Eur J Clin Nutr 2001; 55: 959–66.

43 Michaud A, Vainik U, Garcia-Garcia I, Dagher A. Overlapping neural endophenotypes in addiction and obesity. Front Endocrinol (Lausanne) 2017; 8: 127.

44 Curtain F, Grafenauer S. Plant-Based Meat Substitutes in the Flexitarian Age: An Audit of Products on Supermarket Shelves. Nutrients 2019; 11: 2603.

45 Ley RE, Bäckhed F, Turnbaugh P, Lozupone CA, Knight RD, Gordon JI. Obesity alters gut microbial ecology. Proc Natl Acad Sci 2005; 102: 11070–5.

46 Tonstad S, Butler T, Yan R, Fraser GE. Type of vegetarian diet, body weight, and prevalence of type 2 diabetes. Diabetes Care 2009; 32: 791–6.

47 Biswal BB, Mennes M, Zuo X-N, et al. Toward discovery science of human brain function. Proc Natl Acad Sci 2010; 107: 4734–9.

48 Muga MA, Owili PO, Hsu C-Y, Rau H-H, Chao JCJ. Dietary patterns, gender, and weight status among middle-aged and older adults in Taiwan: a cross-sectional study. BMC Geriatr 2017; 17: 268.

49 Hall KD, Ayuketah A, Brychta R, et al. Ultra-processed diets cause excess calorie intake and weight gain: an inpatient randomized controlled trial of ad libitum food intake. Cell Metab 2019.

50 Forestell CA. Flexitarian Diet and Weight Control: Healthy or Risky Eating Behavior? Front Nutr 2018; 5.

51 Knowler WC, Barrett-Connor E, Fowler SE, et al. Reduction in the incidence of type 2 diabetes with lifestyle intervention or metformin. N Engl J Med 2002; 346: 393–403.

52 Kuna ST, Reboussin DM, Borradaile KE, et al. Long-term effect of weight loss on obstructive sleep apnea severity in obese patients with type 2 diabetes. Sleep 2013; 36: 641–9.

53 Li G, Zhang P, Wang J, et al. Cardiovascular mortality, all-cause mortality, and diabetes incidence after lifestyle intervention for people with impaired glucose tolerance in the Da Qing Diabetes Prevention Study: a 23-year follow-up study. lancet Diabetes Endocrinol 2014; 2: 474–80.

54 Dattilo AM, Kris-Etherton PM. Effects of weight reduction on blood lipids and lipoproteins: a meta-analysis. Am J Clin Nutr 1992; 56: 320–8.

55 Wing RR, Lang W, Wadden TA, et al. Benefits of modest weight loss in improving cardiovascular risk factors in overweight and obese individuals with type 2 diabetes. Diabetes Care 2011; 34: 1481–6.

56 Foster GD, Borradaile KE, Sanders MH, et al. A randomized study on the effect of weight loss on obstructive sleep apnea among obese patients with type 2 diabetes: the Sleep AHEAD study. Arch Intern Med 2009; 169: 1619–26.

57 Warkentin LM, Das D, Majumdar SR, Johnson JA, Padwal RS. The effect of weight loss on health-related quality of life: systematic review and meta-analysis of randomized trials. Obes Rev 2014; 15: 169–82.

58 Schreiner P, Yilmaz B, Rossel J-B, et al. Vegetarian or gluten-free diets in patients with inflammatory bowel disease are associated with lower psychological well-being and a different gut microbiota, but no beneficial effects on the course of the disease. United Eur Gastroenterol J 2019; : 2050640619841249.

59 Pfeiler TM, Egloff B. Examining the “Veggie” personality: Results from a representative German sample. Appetite 2018; 120: 246–55.

60 Cole M, Morgan K. Vegaphobia: derogatory discourses of veganism and the reproduction of speciesism in UK national newspapers 1. Br J Sociol 2011; 62: 134–53.

61 MacInnis CC, Hodson G. It ain’t easy eating greens: Evidence of bias toward vegetarians and vegans from both source and target. Gr Process Intergr Relations 2017; 20: 721–44.

62 Thomas MA. Are vegans the same as vegetarians? The effect of diet on perceptions of masculinity. Appetite 2016; 97: 79–86.

63 Garnett EE, Balmford A, Sandbrook C, Pilling MA, Marteau TM. Impact of increasing vegetarian availability on meal selection and sales in cafeterias. Proc Natl Acad Sci 2019; 116: 20923–9.

64 Johnston BC, Zeraatkar D, Han MA, et al. Unprocessed red meat and processed meat consumption: dietary guideline recommendations from the nutritional recommendations (NutriRECS) consortium. Ann Intern Med 2019.

65 Slanger T, Mutschelknauss E, Kropp S, Braendle W, Flesch-Janys D, Chang-Claude J. Test– retest reliability of self-reported reproductive and lifestyle data in the context of a German case– control study on breast cancer and postmenopausal hormone therapy. Ann Epidemiol 2007; 17: 993–8.

